# The human voice aligns with whole-body kinetics

**DOI:** 10.1101/2023.11.28.568991

**Authors:** Wim Pouw, Werner Raphael, Lara S. Burchardt, Luc P.J. Selen

## Abstract

Humans often vocalize while concurrently gesturing with their hands. Fluctuations in the intensity and tone of the voice have been shown to synchronize with gestural upper limb movement. This research provides direct evidence that interactions between arm movements and postural muscle activity cause these voicing fluctuations. We show that specific muscles (e.g., pectoralis major, erector spinae), associated with upper limb movement and their postural anticipations, are especially likely to interact with the voice. Adding mass to the upper limb increased this interaction. Ground-reaction forces were also found to relate to postural muscles, and these measurements also directly covaried with fluctuations in the voice during some movement conditions. These results show that the voice co-patterns with whole-body kinetics, i.e. forces. We thereby go beyond kinematic analyses in studying interactions between gesturing and vocalization, invoking several implications for biomechanical modeling. We conclude that human voicing has evolved in a dynamical interaction with the whole-body motor system.

---

In textbooks about the mechanics of respiration, vocalization, and speaking, students learn that there are primary and secondary respiratory muscles. The intercostal muscles and diaphragm are taught to be the “primary” respiratory muscles that control subglottal pressures needed to phonate at normal levels, whereas the “secondary” or “accessory” muscles, like the abs or muscles in the back, are only recruited under extreme conditions of breathing and/or speaking (e.g., loud speech, shouting; (1)). Yet these “secondary” muscles are actually very often recruited during speaking because humans often produce hand-gestures, upper limb movements that are communicatively significant (2–4). Namely, humans often produce beat gestures, which are pulsing arm movements that will activate trunk muscles (3, 5). These trunk muscles are known to support forced expiration during more extreme respiratory situations such as coughing (6). Furthermore, muscles around the shoulder girdle are active during upper-limb gesturing that accompanies speech (7, 8). While these muscles are not often activated for speech or tidal breathing alone (1), they are known to be used for respiration in persons with respiratory problems such as COPD (9, 10). Given that these muscles *are* active during (gestural) arm movements that accompany speech in typical able-bodied humans (2, 3, 11), they might regularly enter into functional synergies with respiration for speaking (similar to people with COPD). The contradiction is that when we speak at normal sound levels, we gesture and therefore activate muscles that are usually described as only activated in extreme respiratory(-vocal) conditions, such as coughing or shouting. This contradiction highlights that a scientific understanding of the respiratory foundations of speaking (12) requires a reconsideration of speaking mechanics during *utterances in action*.

Hand gestures are the most common type of action produced during speaking. Gestures often have a pulsing, ‘beat-like’, quality with sudden stops and accelerations that are temporally integrated with vocal aspects of speech (13). These beating movements are found to be strategically timed with excursions in voice intensity and fundamental frequency (F0) and have also been associated with phrase-final lengthening (14). These acoustic features contribute to the perception of so-called accented or stressed moments in speech (13), i.e., they contribute to the multimodal prosody of speech. Multimodal prosody (13) concerns the study of how movement and speech couple, and has been of interest to linguistics (15), but generally ignored in kinesiological research. Traditional linguistics research on multimodal prosody considers the recruitment of gestural movements as salient *visible* signals, synchronized with speech through cognitive mediation (13). Here we hypothesize that the underlying gesture’s movement *dynamics* cause changes in the acoustic features of the voice. In other words, we argue that gesturing and vocalization are synchronized through whole body *kinetics*.

It is only recently that gesture-speech synchronization has been theorized to have its origins in biomechanics (16). The available evidence for the biomechanics thesis is from kinematic research and not from research monitoring muscle activity directly (17–25). Namely, more extreme peaks in the acceleration of movements with larger (arm) vs. smaller (wrist) upper limb segments relate to larger chest-circumference changes, which is found to relate to more extreme acoustic effects on the intensity of vocal sound (21). Furthermore, acoustic effects of upper limb movements are more extreme when subjects are in a more unstable standing versus sitting position (26). These previous studies assessed continuous voicing (19, 26), mono-syllable utterances (21), singing (18, 27), and fluent speech production (20). The combined evidence supports the gesture-speech biomechanics thesis that a physical impulse *J* (mass x acceleration integrated over some time window) impacts posture (especially when standing) and recruits muscles around the chest, which interact with or recruit respiratory-related muscles that affect rib cage movements, as such modulating respiratory-vocal functioning (such that intensity and secondarily F0 are affected). The physical impulse needs to be of a certain still unknown threshold however, as it is clear that movement effects on vocalization diminish when lower mass movements are involved (19–21, 26), or diminish when accelerations are lower (18, 20), or disappear completely, when accelerations close to constant during low-resistance arm cycling (24).

According to the gesture-speech physics hypothesis (16), the effects of upper limb kinematics on vocal utterances lie in biomechanical interactions. However, the studies relating to this theory have so far not been explicitly performed at the level of these *kinetic* interactions (17–22, 24–26, 28, 29). The hypothesis is that physical impulses, that relate to the mass and the acceleration of the upper limb segment, recruit focal and/or more peripheral postural muscles that are also involved in respiratory-vocal control (2, 3, 30). For example, the pectoralis major drives internal rotation of the upper arm and stabilizes other arm movements, but is also associated with forced respiratory action (1, 31). However, most upper limb actions involve a coordinated activation of a whole suit of muscle units around the trunk and shoulder girdle, acting as a pre-stressed ‘tensegrity’ system (32). This tensegrity structure anticipates and reacts to the destabilizing effects of movement on body posture (33, 34). Previous research indeed indirectly indicates possible interactions: the erector spinae is considered a postural muscle during upper limb movement (11), but due to its attachment to the rib cage it is also implicated in “secondary” respiratory control (1). In non-human animals, trunk muscle integration into respiratory action has been well-documented. For example, in flying echo-locating bats (*Pteronotus parnellii*; (35)) muscles activated for flying (pectoralis) are at the same time mechanically driving expiration for echo-vocalizing. However during passive vocal echo-locating, muscle units around the abdomen (rectus abdominis) will drive expiration. This biomechanics research suggests that different muscle synergies are responsible for the expiratory vocal flow of vocalization during flight versus passive activities (see also similar evidence of synergies in courtship displays: (36)).

We hypothesize that also in humans, movements of the upper limbs are biomechanically integrated with vocalizations. However, no direct evidence has been provided whether a particular type of muscle activity, postural destabilization, and mass (rather than size) of the body segment set in motion, predicts inflections in vocalization. Therefore we must go beyond kinematic measures and assess signatures of bodily *kinetics*, the posture-perturbing muscle-generated physical impulses. These impulses affect the ‘tensegrity’ – naturally pre-stressed - musculo-skeletal system (37), which distribute forces over tensioned muscle chains that affect respiratory-vocal functioning, making the voice momentarily louder (a key prosodic feature of the voice). We show that the voice aligns with whole-body kinetics, measuring activity from several muscles, postural perturbations and concomitant voice modulations.

## Current study and hypotheses

In this study (see supporting methods for details, see Figure 1 for a schematic overview) we measured the voice acoustics of participants vocalizing a sustained /a/ vowel (<u>sound</u> <u>examples</u>). Upon 3 seconds of vocalization, participants were prompted to make a single upper-limb movement while keeping their vocalization as steady as possible for another 4 seconds. Participants made different upper limb movements (extension, flexion, internal rotation, external rotation, no movement), with and without a 1kg wrist-weight, while we measured muscle activity (surface EMG) and concurrent postural perturbation (vertical ground-reaction measurements). Figure 1 provides an overview of the experiment, and an example of time series data collected.

**Figure 1.**
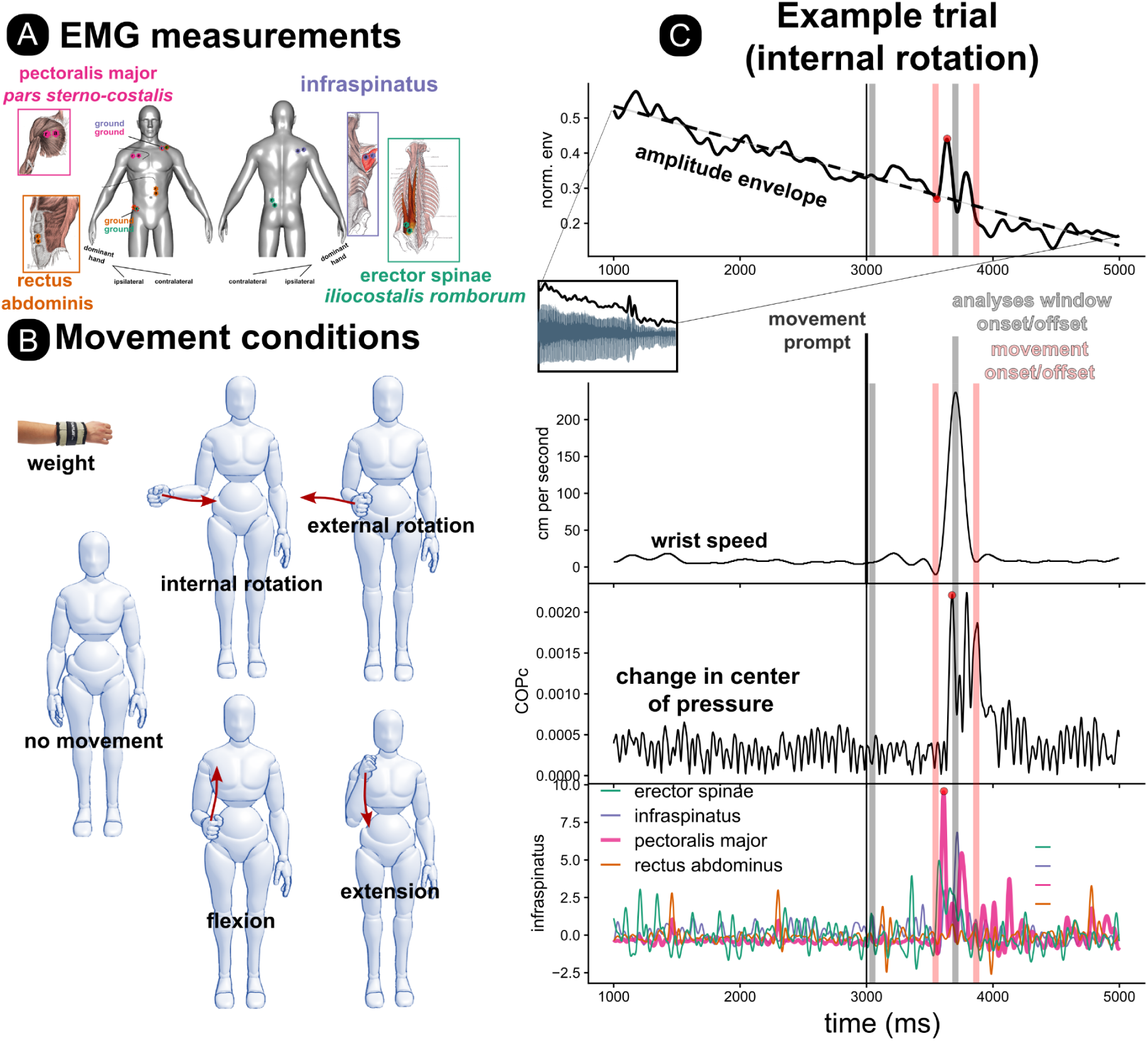
Experimental methods. A) During the experiment we collected surface ElectroMyoGraphy (sEMG) to track muscle activity at four locations: pectoralis major [internal rotator], infraspinatus [external rotator], erector spinae [postural], rectus abdominis [postural]. B) Participants performed vocalizations while wearing a wrist-weight or no weight, and performing several movement conditions. C) For each vocalization trial (example of an internal rotation trial shown here), lasting 7 seconds with a movement prompt presented on the screen after 3 seconds (here 4 seconds is shown), a smoothed amplitude envelope was extracted (z-normalized units, normalized per participant), and a linear trend line was determined to express negative and positive vocal peaks relative to this trendline. The vocal acoustics, muscle activity and postural peaks (global maxima) were determined between 500 milliseconds before movement onset and the moment of peak speed (this ensures to only consider effects of movement onset, rather than the moment of deceleration). Peak speed is used because it captures the moment between acceleration phase and the deceleration phase (and it has one global maximum peak; see text for further explanation).

We made the following predictions with accompanying pre-registered analysis that should help us assess the overall research question of whether kinetic aspects of upper limb movement affect vocalization through (posture-related) muscle activations:

I. Do (different) movement (and weight) conditions have (different) effects on vocalization amplitude? I Null hypothesis H0: The movement and weight conditions, as compared to control condition (no movement, no weight condition) does not affect vocalization amplitude.
II. II. Is muscle activity (differentially) affecting the amplitude of vocalization? II Null hypothesis H0 : Muscle activity induced by limb movement does not relate to vocalization amplitude in any of the conditions.
III. Is specific muscle activity related to postural stability? III Null hypothesis H0: Muscle activity does not relate to postural stability in any of the conditions.

If arm movement affects vocalization as compared to passive condition, and if adding a 1kg weight increases this interaction with vocalization, and if muscle activity is directly related to vocalization, then these overall analyses strongly support the kinetic connection of the movement-vocalization synchrony. Additionally, if specific muscles (rectus abdominis, or erector spinae) are reliably affecting the voice and are also implicated in postural stability, then we have a good basis for concluding that some muscles involved in postural stability are contributing to arm-movement effects on vocalization, a finding we only indirectly observed in previous research showing that gesture-vocal interactions increase in a standing versus sitting position (26). We will also perform several exploratory analyses that help qualify the main analyses (e.g., timing relations between postural, kinematic, muscle, and voice, changes). For the pre-registered analyses we focused on two markers of the voice, namely positive peaks in the amplitude envelope, which can inform about increased expiratory pressures (compressive effects on the lungs), and negative peaks, which would indicate inspiratory pressure (expansive effects on the lungs).

## Methods

Informed by a simulation-based power analysis for the three confirmatory analyses (38) reported in the pre-registration (https://osf.io/jhdq4^1^) we recruited *N* = 17 participants: 7 female, 10 male, *M* (SD) age = 28.50 (6.50), *M* (*SD*) body weight = 72.10 kg (10.20). Note that our digital Supplemental Information with reproducible code is provided on our <u>Github webpage</u>.

The study consisted of a 5-level within subject design’ of movement condition: <u>no</u> <u>movement</u>, as well as four actions with the elbow flexed at a 90-degree angle: <u>flexion</u> and <u>extension</u>, <u>internal rotation</u>, and <u>external rotation</u> (see Figure 1); these latter four conditions we will refer to as the “movement-only conditions”. Additionally, there was a two-level within subject weight condition, where participants performed movements with and without a 1kg wrist weight. Weight was manipulated to increase the physical movement impulse. Finally, there was a 2-level within-subject vocal condition where participants either expired versus vocalized during the trial. To reduce the number of comparisons, and as stated in our pre-registration, we only focus on the vocalization condition in this confirmatory research, and only take into account the expiration condition for some exploratory confound analyses. Thus we are left with a 5 (Movement condition) x 2 (Weight condition) fully within-subject design. We were able to collect 1281 experimental trials, with 5% of trials that were not recorded due to experiment software failure (17 participants x 5 movement conditions x 2 weight conditions x 2 vocal conditions x 4 repetitions = 1360), entailing 2.7 hours of continuous voice data. For the vocal condition, which is our main focus here, we collected 636 trials (17 participants x 5 movement conditions x 2 weight conditions x 2 = 680, minus 6.5% unrecorded trials with software issues).

### Alpha correction

To correct for multiple testing, we apply a Bonferroni correction on the alpha level (i.e. the significance level). We perform three tests, so the new alpha level necessary to reject the null hypothesis is .016 (.05/3=.016). This alpha level is also adopted for exploratory findings.

### Analysis procedure

Our analysis is aimed to relate vocal inflections with postural and muscle activity due to upper limb movement. We therefore identified positive and negative peaks relative to a trend line (see below) in the smoothed amplitude envelope of the vocalization before the peak in wrist speed was reached (see <u>Supplemental Methods</u> for further detail). Thereby we assess peaks in vocalization related to the initiation of the movement rather than the physical impulse produced during deceleration (which will recruit antagonistic muscle units as compared to the initiation phase). Note that there is a general decline in the amplitude envelope of the vocalization during a trial (see Figure 1) as the subglottal pressure falls when the lungs deflate. To quantify deviations from stable vocalizations, we therefore detrend the amplitude envelope time series, to assess positive or negative peaks *relative to this trend line*. For the envelope, muscle activity, and the change in center of pressure we measure the global maxima happening within the analyses window (i.e., within a trial we take a local maximum occuring between movement onset and offset). We analyze positive and negative peaks within the movement window separately. To ensure that we identify multimodal signal peaks that are associated with the onset of a movement we determine the peak speed of a movement and look for multimodal signal peaks (vocal, EMG, ground reactions) before that peak in speed. Peak speed is used because this allows us to identify the moment before the deceleration starts to take place (i.e., the moment where the acceleration = 0 cm/s^2^). A deceleration entails a new muscle synergy that needs to break the moving body segment (which would complexify analyses if taken into account in the current analyses). Peak speed is a more convenient anchor because it is the global maximum of the speed time series in the trial. We replicate the analysis procedure for the no movement condition as a control comparison, though effectively we are identifying random moments during the vocalization as there is no significant movement speed peak.

### Descriptives and manipulation checks

Before assessing the main hypotheses, initial information about the timing of the different modalities as well as manipulation checks on how the condition affects muscle activations is informative.

### Manipulation Check: Muscle and postural activity for movement vs. no movement condition

Figure 2 provides an overview of the magnitude of the peaks for muscle activity, changes in center of pressure, and wrist speed (see Table S1 and Figure S4 for more complete numerical information). It is apparent that depending on whether there is a movement or not, there is more muscle and postural activity. To assess this statistically, we ran a mixed regression model with participant as random intercept and movement vs. no movement as independent factor, and assessed whether this was a better predicting model than a base model predicting the overall mean for the different muscle activities. All muscles (change in **χ**^2^ (1) > 113.30, *p*’s < .0001), except rectus abdominis (change in **χ**^2^ (1) = 3.78, *p* = .0518) were better predicted by movement vs. no movement as compared to a base model. Movement vs. no movement also predicted changes in the center of pressure (in any direction) better than a base model (change in **χ**^2^ (1) = 360.90, *p* < .0001).

**Figure 2.**
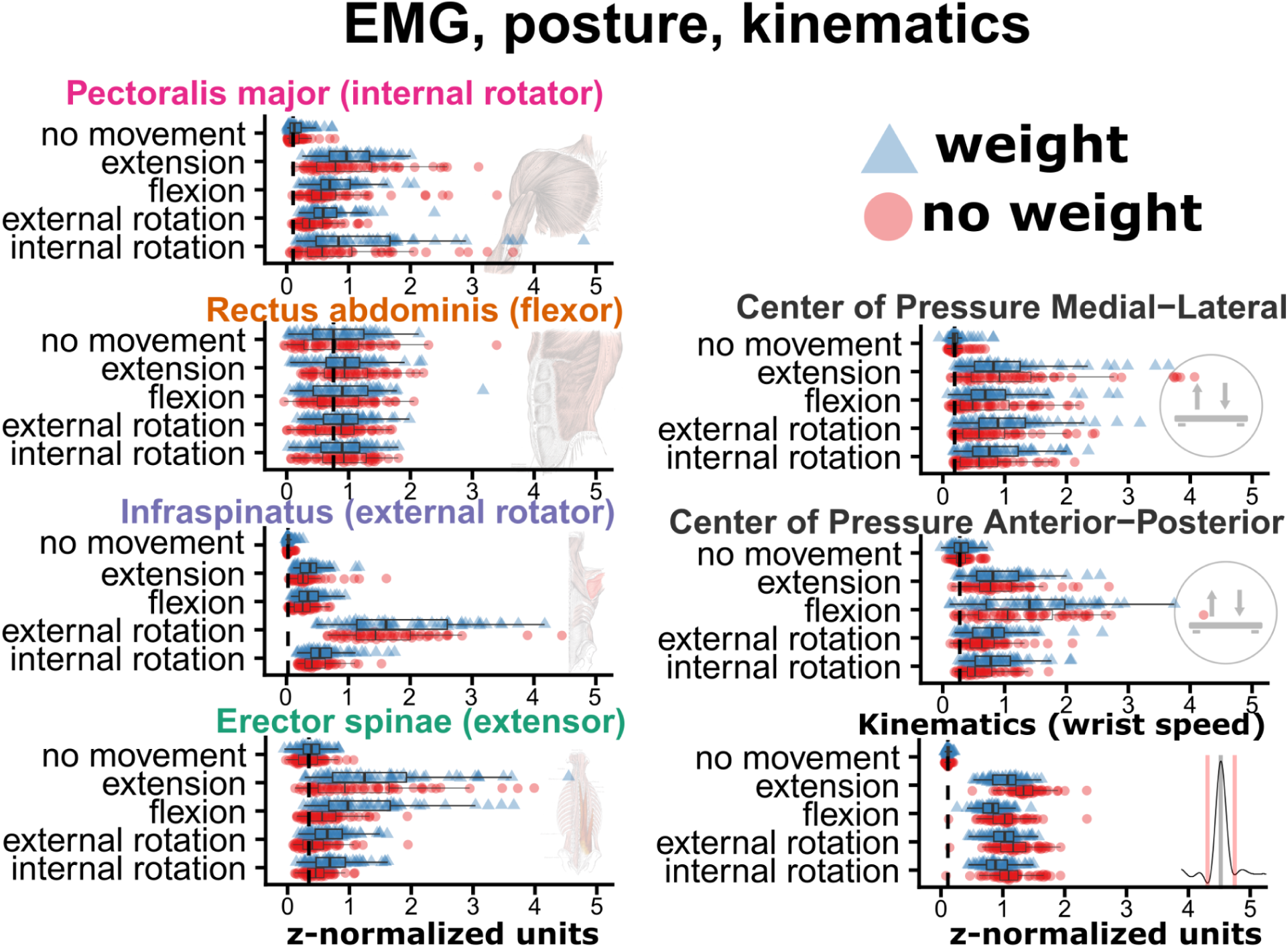
Manipulation checks of changes in muscle, posture, and kinematics, based on movement and weight condition. It is clear in C) that movement coincides with peaks in muscle (especially pectoralis major), postural, vocal acoustics. D) provides the muscle and postural activity (z-normalized units, normalized by participant).

### Manipulation Check: Muscle and postural activity, and kinematics for different weight conditions

In Figure 2, it is clear that in general there is an increase in muscle and postural activity in the weight condition. We assessed whether the presence or absence of a 1kg weight during trials in the movement condition affected muscle activity over and above any effect of the different types of movements. The activity of all muscles (change in **χ**^2^’s (1) >15.68, *p*’s < .0001), except rectus abdominis (change in **χ**^2^ (1) = 0.08, *p* = .7721) was better predicted by an independent factor for weight vs. no weight as compared to a base model predicting the overall mean. The added effect of weight was also present when predicting changes in center of pressure (in any direction, in **χ**^2^’s (1) = 1.26, *p* = .2614).

In Figure 3, it can be seen that the wrist speed seems to decrease when wearing a wrist weight, and in the supplemental materials Figure S4 it is also clear that wrist acceleration decreases. Indeed, a model with a factor for wrist weight reliably predicted peak speed (change in **χ**^2^ (1) = 204.43, *p* < .0001) and peak acceleration (change in **χ**^2^ (1) = 257.77, *p*’s < .0001) better than a model with movement condition as only predictor. We can therefore conclude that adding a wrist weight does slow down movement, but it also recruits higher muscle activity (except for the rectus abdominis), suggesting an overall increase in exerted effort. Given the decrease in acceleration, we must conclude that the potential effect of increased mass on the kinetics (physical impulse) must be reduced as compared to when kinematics remain unchanged.

**Figure 3.**
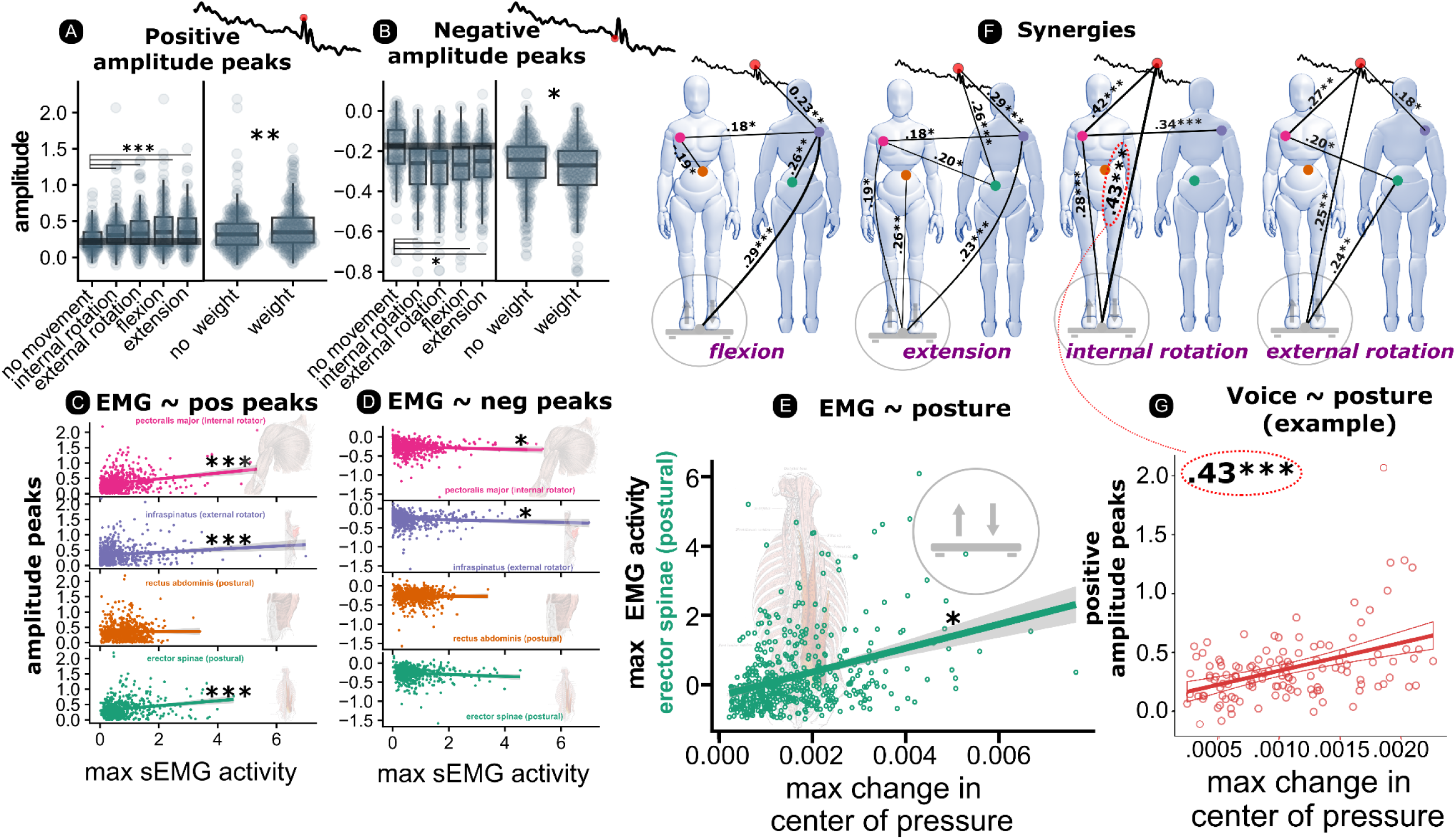
Overview Main results. A) shows the magnitude of positive peaks (relative to trendline) for the different movement conditions. B) shows this for the negative peaks (note A and B have different y-axis ranges). Plots C and D show the relationship between peak muscle activity and positive (C) and negative (D) vocal amplitude envelope peaks. E) shows the relationship with the change in center of pressure and the peak activity in the postural muscle erector spinae. All axes are given in z-normalized units (normalized within participants). F) provides the statistically reliable Pearson correlation coefficients (line thickness also represents correlation strength) between all the measurements (max EMG pectoralis major in pink, infraspinatus in purple, erector spinae in green, positive peak amplitude envelope in red, and the change in center of pressure in grey), presented separately for movement condition. G) highlights the highly reliable correlation obtained in the internal rotation condition (in F) between change in the center of pressure and the magnitude of the positive peak in vocal amplitude envelope (i.e., posture directly relates to voice). * p < 0.05; ** p < 0.01; *** p < .001

### Manipulation Check: Differentiated effects on muscle and postural activity for different types of movement

In Figure 3, it seems apparent that for each movement condition there is a differentiated recruitment of the muscles, and different patterns of anterior-posterior and medial-lateral changes in center of pressure. We see for example in Figure 3 that the infraspinatus has much higher activity in the external rotation condition, while the pectoralis major has much higher activity in the internal rotation condition. From Figure 3 we can also see that for the anterior-posterior movements (flexion and extension) there is generally higher activity for the erector spinae and changes in anterior-posterior changes in the center of pressure, which would mean that not only focal muscles, but also postural muscles, are differentially affected. We confirmed that there was indeed a differentiated effect of movement type on the muscles by again comparing a null model predicting the overall mean against a model with movement conditions (excluding no movement, making this a conservative test) as a factor. We found that all muscles were differentially affected by movement condition (change in **χ**^2^ (3) > 107.85, *p*’s < .0001), except the rectus abdominis (change in **χ**^2^ (3) = 2.20, *p* = .5312). We also found that posture activity in the anterior-posterior (change in **χ**^2^ (3) = 433.63, *p* < .0001) and the medial-lateral direction (change in **χ**^2^ (3) = 43.10, *p* < .0001) was differentially affected by movement condition.

### Summary manipulations checks

● Each muscle, except rectus abdominis, increases in peak EMG magnitude during movement vs. no movement condition
● Each muscle, except rectus abdominis, has (some) differences in peak EMG magnitude levels per movement type
● Each muscle, except rectus abdominis, increases in peak EMG magnitude in the weight vs. no weight condition
● Peak magnitude in center of pressure (overall) increases during movement vs. no movement condition
● Peak magnitude in center of pressure (anterior-posterior) has different levels per movement type
● Peak magnitude in center of pressure (medial-lateral) levels per movement type
● Peak magnitude in center of pressure (overall) increases in the weight vs. no weight condition
● Peak magnitude in speed and acceleration, decreases in the weight vs. no weight condition

### Main results

Please see the supporting results for a full report of all the analyses (for a computationally reproducible version of the extended results with code <u>see here</u>). Also, see Figure 3 for an overview of the main results.

### Research Question 1: Do (different) movement (and weight) conditions have (different) effects on vocalization amplitude?

A mixed linear regression model with weight and movement conditions and participant as random intercept explained more variance than a base model predicting the overall mean vocalization amplitude (hange in **χ**^2^ (5) = 36.02, *p* < .001). Compared to vocalizing without a limb movement, movement conditions (internal rotation, external rotation, flexion, extension) lead to more extreme positive peaks in the amplitude envelope of the vocalization (*b’s* > 0.116, *SE*’s < 0.032, *p*’s < .001; see Table S7). Also negative vocalization peaks are systematically affected by the weight and movement conditions, as compared to a base model (hange in **χ**^2^ (5) = 48.47, *p* < .001). Movement conditions all increased the magnitude of negative vocal peaks (*b’s* > 0.080, S*E*’s < 0.018, *p*’s < .001; see Table S5). Thus no matter what type of movement, the voice is affected, leading to positive-valued and negative-valued perturbations in the intensity of the voice. Exploratory post-hoc analyses (Tukey corrected; see Table S11) confirmed individual effects of movement types as compared to no movement (*p*’s < .003), but did not reveal differences between the different types of arm movements in terms of vocal effects (*p*’s > .346).

In the movement conditions, weight increases the magnitude of the positive vocalization peaks (*p* = .01; see Table S6), but leaves the negative peaks unaffected (*p* = .024; see Table S6)^2^. Thus by increasing the mass of the upper limb with 1kg, the magnitude of the positive vocalization peaks is also systematically increased.

We therefore reject the null hypothesis for research question 1, and conclude that weight and movement condition increases positive peaks in the amplitude of the voice. Though differential effects between movement types are not observed.

### Research Question 2

Is muscle activity (differentially) affecting the amplitude of vocalization?

We further assessed how different general muscle activity relates to positive and negative peaks in the vocalization acoustics in an analysis where we pool all data (disregarding movement condition). We performed a linear mixed regression (with participant as random intercept) with a model containing peak EMG activity for each muscle which was regressed on the positive peaks in vocalizations. We obtained that a base model predicting the overall mean of positive peaks was outperformed relative to said model (change in **χ**^2^ (4) = 59.08, p < .001). Positive peaks in vocalizations were related to activity in pectoralis major (*b* = 0.013, *SE* = 0.003, *p* < .001), erector spinae (*b* = 0.016, *SE* = 0.004, *p* < .001), and infraspinatus (*b* = 0.006, *SE* = 0.002, *p* = .005), but not in rectus abdominis (*b* = -0.003, *SE* = 0.007, *p* = .710), as shown in Table S7. In general, we obtained more noisy measurements in the rectus abdominis activity, also having to do with there being more adipose tissue. None of the muscles were reliably associated (at *p* < .016) with negative peaks in vocalization and the relations were weaker than the comparison with positive peaks (*b*’s < .004, SE’s < .002, *p’s* < .020), though the general model comparison against a base model did reach statistical significance (change in **χ**^2^ (4) =16.54, *p* < .001).

We therefore reject the null hypothesis for research question 2, and conclude that focal and postural muscle activity is related to positive peaks in the amplitude of the voice.

### Research Question 3: Is postural muscle activity related to postural stability (change in center of mass)?

The peak in the change in the center of pressure with muscle activity was indeed related to postural muscle activity. We obtained that a base model predicting the overall mean of the change in the center of pressure (COPc) was outperformed relative to a model containing peak EMG activity for each muscle, Change in **χ**^2^ = 180.94 (4), *p* < .001. Namely, the erector spinae (*b* = 0.133, SE = 0.012, *p* < .001) had the highest association with change in the center of pressure. However, we also found that pectoralis major (*b* = 0.051, SE = 0.009, *p* < .001) was related to postural changes, as shown in Table S14. The rectus abdominis and infraspinatus did not show this relationship in a pooled analysis (*b’s* < 0.034, SE’s > 0.012, *p* > .134).

We therefore reject the null hypothesis for research question 3, and conclude that postural muscle activity (erector spinae) is related to changes in center of pressure which index postural instability.

### Exploratory analyses: Synergies

For the previous analyses, we lumped all movement conditions together to assess relationships between muscle activity, posture, and vocalization. However, we also investigated context-dependent, i.e. movement condition, variability in the dependent variables. We computed Pearson correlation matrices between the positive peaks in EMG, change in COP and vocalization amplitude for the individual movement conditions. Separating out, rather than pooling our analyses over the movement conditions generates some interesting patterns. For example, the rectus abdominis has generally been observed as non-responsive in our different analyses, but in the extension condition we do implicate rectus abdominis to postural control (Pearson *r* = .26, *p* < .01; see Table S17). We further obtain a very strong association in the internal rotation condition for the degree to which the body changes center of pressure and the magnitude of the positive peak in vocalization (see Figure 3, panel F & G), Pearson *r* = .43, *p* < .001 (see Table S10 or Figure S9). This shows that posture perturbations can be directly related to perturbations in the vocalization. In general, it seems that the pectoralis major is a key muscle that relates to the voice, or it relates to another muscle activation that relates to the voice, across all movement types.

### Exploring potential confounds: Causality direction

It is quite possible that effects on vocalization are fully determined by the focal muscles that move the arm. Anticipatory postural muscles then correlate with that focal activity, without contributing to effects on the voice. However, we think depending on the type of movement, a combination of different muscles that affect movements of the surrounding structures of the lungs can contribute to voice modulation.

That there is a complex *combination* of causal relations possible is suggested from additional exploratory analyses reported in Figure 4. Here we have modeled the by-participant normalized (z-scale) continuous trajectories of the amplitude envelope and all the other signals (EMG’s, posture) using Generalized Additive Modeling (GAM) (39) for each movement condition, setting participants as random intercept. Here we ignore the rectus abdominis which has proven rather noisy and unresponsive to our manipulations (but we do show this GAM curve in the supplemental information). To reduce over-estimation of any effects, we also accounted in the model for the autocorrelation of the residuals (24, 39). Based on the predicted non-linear slopes that we generated for an interval of 600 ms we determine firstly the approximated onsets of the continuous measures. Onsets are determined as the peak in the 2nd derivative for time of the GAM trajectory for a particular signal and condition. In general, we only consider the presence of onsets when the overall peak of the signal exceeds 0.75 x standard deviation from the mean (otherwise the timings reported are difficult to compare). For the figure, we thus center our data based on the predicted *onset* that leads to the positive peak in the amplitude envelope. In this way we can isolate when the amplitude envelope starts to respond relative to the other signals.

**Figure 4.**
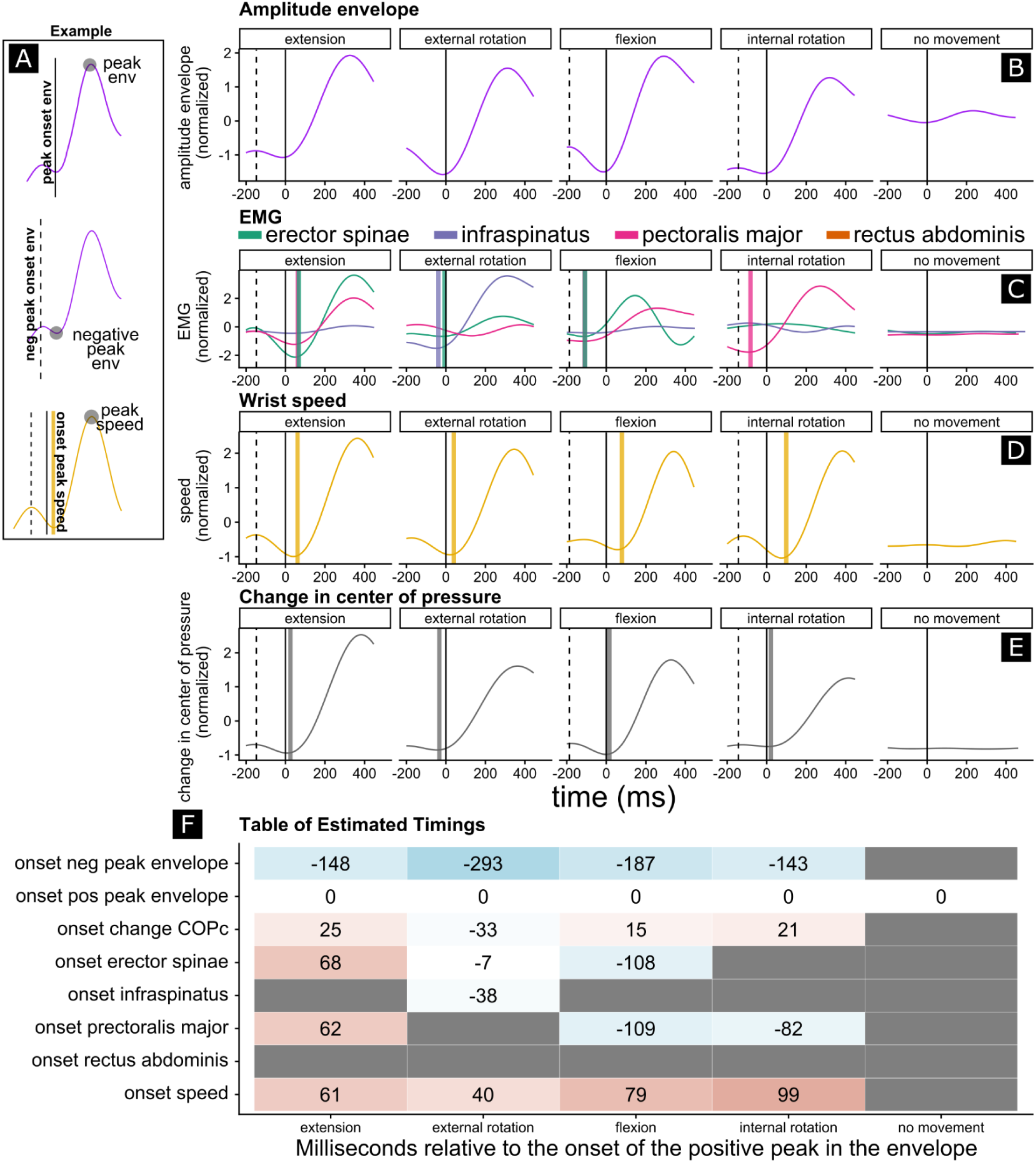
Timing of muscle, postural, and kinematic peaks. As exemplified in Panel A each black vertical line shows the onset towards a negative peak (dashed vertical line) and the onset towards a positive peak (solid vertical line) in the amplitude envelope, which is projected over each subplot that belongs to each modality (B, C, D, E). For the other modalities, a solid vertical thick line is shown for the onset towards a positive peak (e.g., onset towards a positive peak in center of pressure in grey, panel E). It can be seen that the moment of first onset of the peak change in pressure coincides closely to the moment of onset towards the positive peak in vocalization. Panel F, summarizes the timings of these onsets relative to the positive peak in the amplitude envelope, showing for example that in the flexion condition the erector spinae activates 103 milliseconds before the onset of the positive peak in the amplitude. In the internal rotation condition, the pectoralis major activates earliest, with a 86 millisecond lead to the estimated onset of the amplitude envelope to its positive peak. Note in general, that changes in vocalization are estimated to happen before observable changes in movement. Changes in vocalization are sometimes preceded by muscle activity and center of pressure changes. This yields further support that it is forces (kinetics), rather than movement (kinematics), that affect the voice.

Firstly, we determined from the GAM trajectories the estimated onset of the vocalization peak (i.e., the moment at which the amplitude starts to rise to peak envelope), as well as the onset of the approximated negative peak, if present (i.e., the moment at which the amplitude starts to fall to the negative peak envelope). We learn from this that the minor negative peaks tend to precede the positive peak, suggesting a possible ‘anticipatory vocal adjustment’. Importantly, however, in our supplemental materials, we further show that the observed negative peaks do not happen as consistently as compared to the positive peaks (see Figure S9). This further corroborates the idea that positive peaks are directly related to physical impulses, while negative peaks are a type of correlated anticipatory/reactionary activity of the vocal system.

Crucially, we find that in general before movement is observed, as indexed by the onset of the peak in wrist speed, there is a change in the center of pressure. Furthermore, muscle activity of postural muscles, such as the erector spinae during flexion conditions, can be observed even 108 milliseconds before the vocal onset, in line with research on anticipatory postural adjustments (APAs; 2, 4, 30). Even in cases where there are no clear activations of the postural muscles, e.g., as in the internal rotation, it can be seen that the focal muscles (e.g., pectoralis) start to activate 82 milliseconds before the peak in vocal onset. Given that there is both postural and focal muscular activity that precedes the vocal onset slightly, it confirms that voice effects are not simply a cognitive anticipation of a physical impulse that happens later.

### Exploring potential confounds: Subtle differential effects of movement-only conditions on vocalization

From our previous confirmatory peak analyses, we concluded that there were no reliable differences between the different movement conditions in terms of affecting the magnitude of the positive peak in the amplitude of the vocalization. Now that we have modeled the trajectories with GAMs, we can perform a statistical test for whether the different movement conditions affect the vocalization amplitude envelopes *in terms of their trajectory*, rather than simply differing in the magnitude of the peak of the trajectory. We determine the trajectory along an interval of -500ms to +500ms relative to the onset of the arm movement (time = 0 when the movement initiates). The GAM analysis (see Figure 5) is much more powerful as we take in millions of time series observations (as compared to the by-trial peak generation).

**Figure 5.**
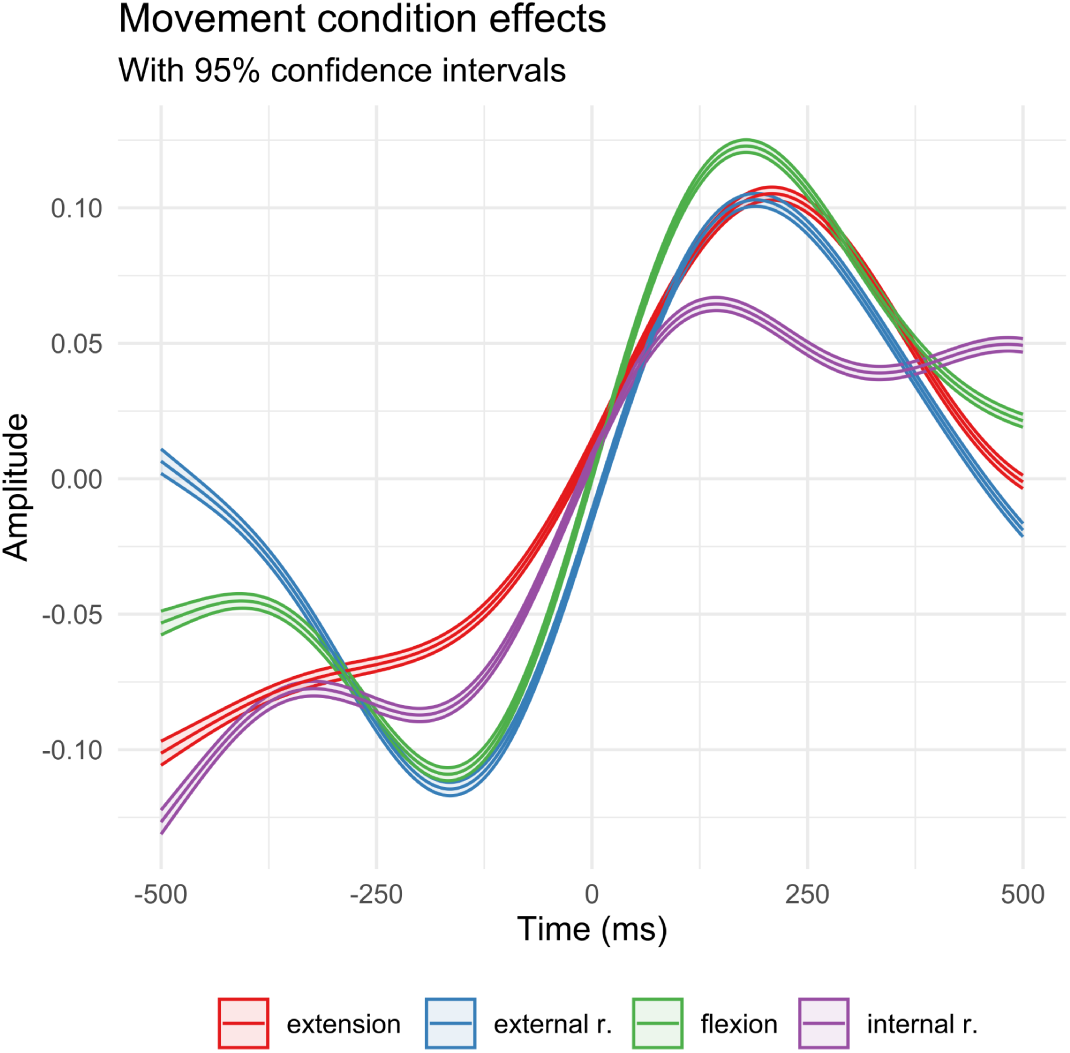
GAM trajectories with 95% confidence intervals for the vocal amplitude envelope for the movement-only conditions (z-scaled units). All trajectories (except extension) showed a movement-specific time course as indicated by the statistically reliable smooths (see Table 1). The trajectories are aligned with the onset of the arm movement (time = 0).

**Table 1.**
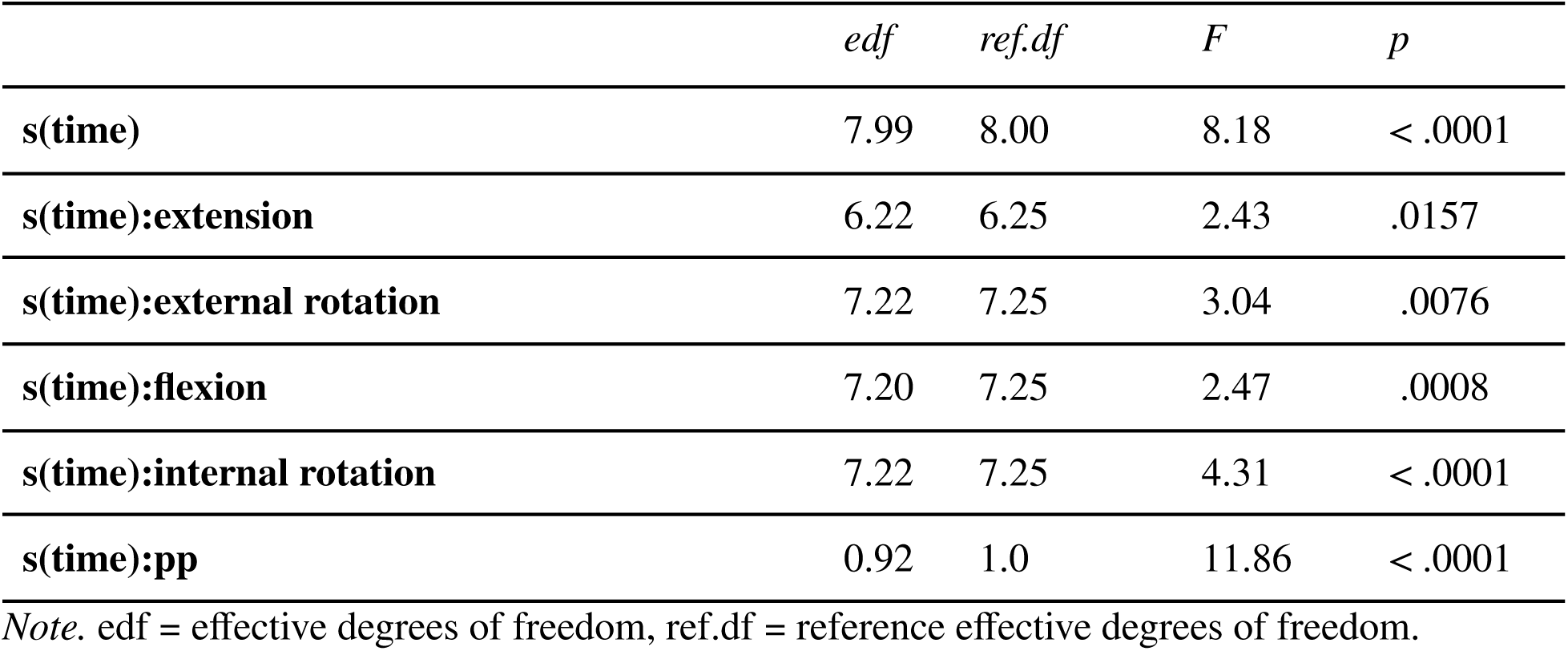
GAM statistics for the trajectories shown in Fig 5.

|  | *edf* | *ref.df* | *F* | *p* |
| --- | --- | --- | --- | --- |
| **s(time)** | 7.99 | 8.00 | 8.18 | < .0001 |
| **s(time):extension** | 6.22 | 6.25 | 2.43 | .0157 |
| **s(time):external rotation** | 7.22 | 7.25 | 3.04 | .0076 |
| **s(time):flexion** | 7.20 | 7.25 | 2.47 | .0008 |
| **s(time):internal rotation** | 7.22 | 7.25 | 4.31 | < .0001 |
| **s(time):pp** | 0.92 | 1.0 | 11.86 | < .0001 |

*Note.* edf = effective degrees of freedom, ref.df = reference effective degrees of freedom.

The GAM model with an autoregressive component, with participants as random intercepts and movement-only conditions, was statistically reliable, *F* (1818052.92, 1818025.83) = 170.97, *p* < .0001. The smooth term statistics in Table 1 show that there was a general nonlinearity to the amplitude envelope that characterized all movement-only conditions, as evidenced by the statistically reliable smooth for time. Additionally, the flexion, external rotation, and internal rotation,each had a movement-specific pattern that was statistically reliable. This means that there are statistically robust deviations in the shaping of the amplitude envelope trajectories for different types of movement (except for the extension condition).

### Exploring potential confounds: assessing overall muscle tension

We explicitly planned in our pre-registration to ignore the expiration condition to decrease the number of analyses reported in the current paper. However, the expiration trials can inform about an important issue that may confound the main results. While we confirmed that there is a differential effect of movements on focal muscles as well as postural muscles, this might be caused by an overall tensioning of the body, just prior to the limb movement, when vocalizing to keep the vocalizations stable. Undifferentiated tensioning of all muscles during vocalization trials may then be a strategy to reduce possible perturbations due to an upper limb movement, thereby inadvertently increasing the likelihood of a perturbation due to increasing postural stiffness which will increase the perturbation strength of the arm-related physical impulses. Thus, under this hypothesis, effects on the voice arise because there is an overall stiffening of the body in anticipation of a very predictable, but - otherwise negligible, physical perturbation from the arm movement. This would reduce the generalizability of our findings as the current effects can only be generated in experimental environments with highly predictable voice perturbations.

In previous exploratory work, we have already established that arm movements also affect acoustic markers of expiratory flow (28), which already weakens the possibility of the confound of keeping vocalization stable. However, here we also test for whether muscle activity was predicted by whether movements were performed under expiration versus vocalizing (over and above the different movement and weight conditions). Activity in the pectoralis major, the infraspinatus, and the rectus abdominis, were not reliably better predicted by vocal condition, over and above weight and movement condition effects, change in **χ**^2^ (1) < 2.594, *p*’s > .107. There was however an overall increased muscle activity over and above weight and movement conditions in the erector spinae when vocalizing (change in **χ**^2^ (1) = 11.797, *p*’s = .0006). Figure 6 provides a descriptive overview of the different muscle activation levels per vocal and movement condition.

**Figure 6.**
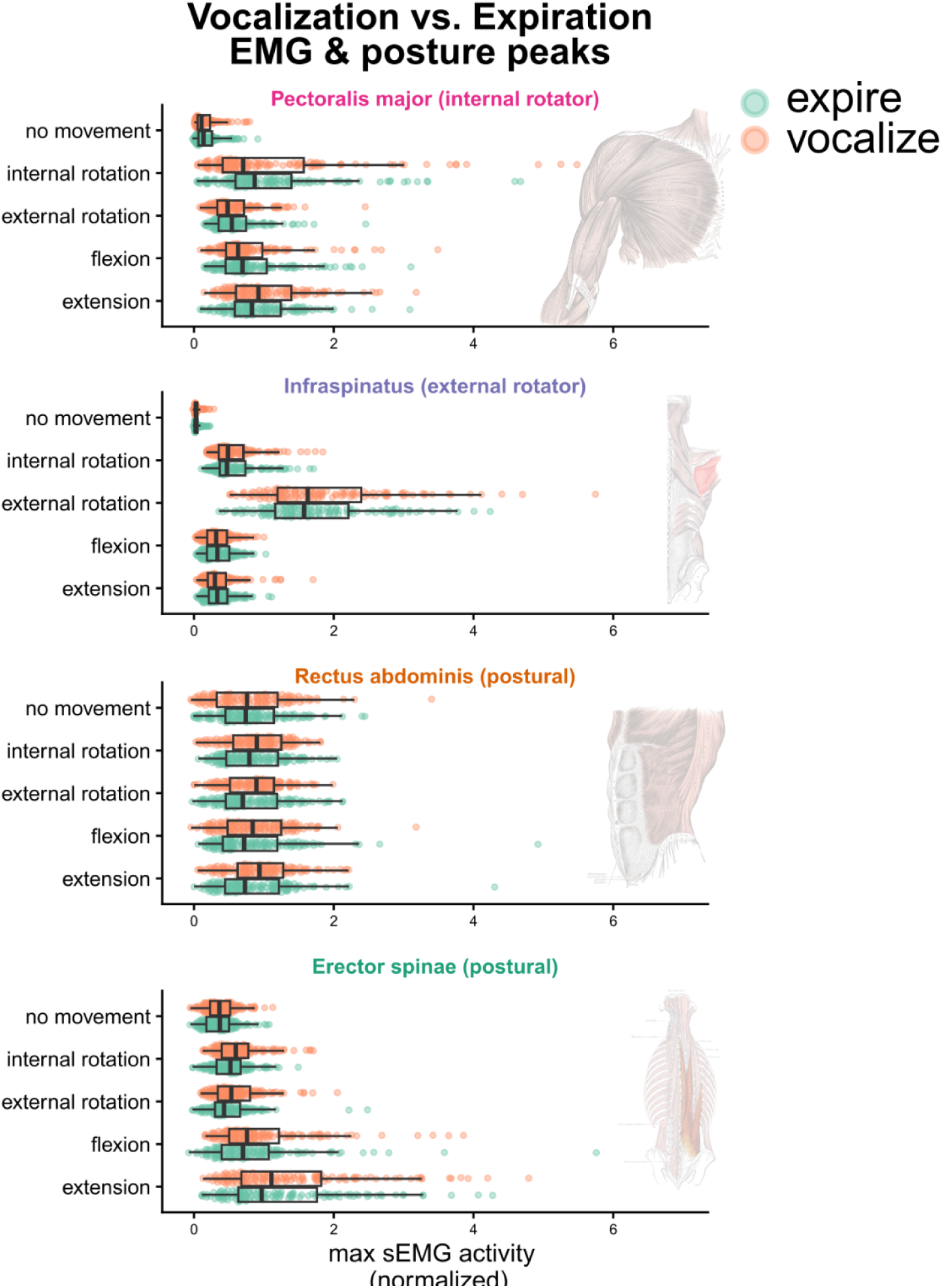
Expiration vs. vocalization. For each vocal condition (vocalization vs. expiration) we show the peak magnitudes of muscle activity per muscle site.

Importantly, it is possible that vocalizing recruits more muscle activity in general, including in the no movement condition, because the subglottal pressures are higher for vocalization due to increased air resistance of the adducted vocal folds as compared to the partially closed lips for controlled expiration. To test an overall increased erector spinae activity, rather than a differentiated increase for movement trials only, we firstly assessed whether a model predicting erector spinae activity for both movement and no movement trials, over and above movement presence and weight conditions, was a better model than a comparison model with only movement and weight condition. This model increased explained variance, change in **χ**^2^ (1) = 9.94, *p* = .0016, and indeed in general there was an overall increased effect of vocalization on erector spinae activity, *b* = 3.152, *t* (3.152) = 0.452, *p* = .0017. Since another model with an added interaction term of movement presence and vocal condition did not reliably improve relative to a model with simple effects, **χ**^2^ (1) = 1.424, *p* = .2326, we conclude that *in general* (also in the no movement control condition) vocalizations recruits higher erector spinae activity than expiration in these tasks. This thereby refutes the possibility that there is increased muscle activation in anticipation of movement perturbing the voice and thereby appearing as if arm movement directly affects the voice.

## Discussion

The current report demonstrates that the human voice resonates with forces produced by upper limb movements. This resonance effect increases if the mass of the limb is increased. Moving the arms requires recruitment of focal (prime mover) muscles, but also postural muscles. Both muscle groups mechanically affect the respiratory system and thus upper limb movements translate into perturbations of the voice’s amplitude when vocalizing. In other words, the voice aligns with whole-body kinetics. We use the term “whole-body kinetics” because not only the focal muscles moving the arm (e.g., pectoralis major), but also trunk muscles stabilizing the body during upper limb movement (erector spinae), are directly related to vocal fluctuations. Further analyses confirm that the activity onsets of focal and postural muscles often precede or coincide with the onset of changes in the vocalization amplitude. Given these timing relations, and given the fact that muscle activation and voice activation relate in magnitude during arm movements, we conclude that both postural and focal muscle activations are likely a cause of the changes in the voice during upper limb movement. Finally, we show subtle differences in the voicing amplitude envelope trajectories across the different movement types. These differences suggest that while there is a net effect of increased subglottal pressures due to arm movements in general, the different movements lead to differently shaped temporal dynamics of the vocalization.

Our confirmatory and exploratory analyses show that it is primarily *positive* rather than negative peaks in the intensity of the voice that are *systematically* arising at particular moments of upper limb movement. It is also positive peaks that are most reliably scaling in magnitude to arm-movement related muscle and postural activity as well as weight condition, suggesting that arm movements primarily cause *increases* in subglottal pressures. This means that during rapid arm movements there is a tendency for (postural) muscles to generate a net compressive effect on the lungs, i.e., a net expiratory effect. This might be because compressive effects on the lungs are functionally enabling the stiffening of the core, serving to increase postural stability. Postural stability is challenged during any upper limb movement (3), and therefore the net counteraction that is needed is a stiffening of the core that has a net expiratory effect. This is a to-be-confirmed hypothesis. Furthermore, in future research we hope to model how particular gradient patterns of muscle activity (within a single type of movement) predicts the shape of vocal trajectory. For example, there will likely be differences over trials or participants in the degree of antagonistic muscle activity of the pectoralis major that stabilizes an external arm rotation - a movement that is primarily driven by the agonistic muscle activity of the infraspinatus. Do these differences in agonist-antagonist activity, and their effects on stiffness, then modulate the perturbing effects of the movement on vocalization? Finally, in our current confirmatory analyses, we have primarily assumed linear relationships between the voice, and muscle and postural activity, but further research can also start taking into account the possible non-linear scaling relationships, which have in several other related studies improved modeling performance (18, 28). By using a linear approach the probability of finding significant effects is reduced compared to non-linear approaches. In sum, further research should investigate in more detail possible context-dependent associations between gesture and vocalization that can show that the voice is shaped by different (non-linear) muscle-voice synergies. The current open data provide a publicly available resource for such investigations (link redacted during review).

There are several further limitations to the current research that should be discussed. Firstly, surface EMG measurements are highly variable in relation to force output, not only because of different measurement conditions for the muscle sites (e.g., differences in adipose tissue across body parts), but also because of individual differences (e.g., differences in adipose tissue across individuals). EMG-force relations also depend on what function the muscles are recruited for (e.g., focal muscles, antagonist stabilizer; (40)). However, our inferences about force and voice are not based solely on the magnitude of EMG activity of the focal muscles (that change functions in the different movement conditions). We paint a much broader picture. Manipulating the mass of the movement segments does not change the function of the moving muscles, but does affect the voice and postural muscles maintain their function under the different limb movement conditions suggesting a direct link between their activity and the observed voice fluctuations. Further support for the latter interpretation is provided by the observed relationship between changes in ground-reaction forces during upper limb movement and voice fluctuations (Figure 3, panel G). Our conclusion that posture is a key part of the voicing dynamics, further resonates with previous research showing that movement-voice coupling is more pronounced in a more posturally unstable standing versus sitting posture (26). This overall picture supports the gesture-speech physics theory (16, 26): gesture-related kinetics result in postural perturbations that affect respiratory-vocal functioning that in part explains gestures’ coupling with vocalization. Note that this does not preclude other processes such as cognitive coordinative stabilities, rhythmic cognition, and language-specific constraints that may bind gesture and speech (16, 41–45).

An additional surprising pattern was that, across the board, we find that the rectus abdominis is largely unresponsive to movement conditions, and does not relate reliably to changes in the voice. We believe that this might be in part a surface EMG measurement issue. Given that the belly area generally involves a higher amount of adipose tissue as compared to the other muscle sites. Therefore, our system likely has not captured this muscle activity well. Needle EMG can provide a better measurement of this muscle, but is rather invasive.

A further challenge of this line of research is how the current findings generalize to more natural speech conditions. There are several important implications of the current results. Firstly, any shortcomings with artificial speech synthesis systems (e.g., models, robots) to reach believably human feature characteristics, need to be aware that disembodied systems are abstracting out how body posture and whole-body muscle chains relate to the voice. Furthermore, our understanding of how effective communication may be compromised by pathologies should be weighed against how the human voice operates in action. It is known that people with Parkinson’s have trouble not only with expressing emotions (46) but also with speech prosody (47, 48), overall appearing monotonic in expression. Obviously, whole-body postural dynamics are compromised in Parkinson’s, which might lead to avoiding effortful gestures, and which then relates to attenuated gesture-vocal communication features.

It has been shown that people who hear another’s vocalization without seeing the vocalizer can glean information about body motions (19, 49). This research indicates that persons are attuning to how the voice can inform about whole-body kinetics, where upper limb physical impulses perturb postural stability, invoking whole-body muscle chains that constrain the respiratory-vocal system. James Gibson (50) in his ecological approach to perception emphasized that the visual system is not limited to the eyes alone. Rather, vision is a dynamic system that involves the entire body and its interaction with the environment. The conclusion is similar when it comes to the voice (51). The human voice is not contained in the larynx. It lies within the tube-like upper vocal tract, lengthened by moving cavities that change resonant properties of the sound (52–54), and it is powered and modulated by respiration, which is situated in a thoracic muscle complex (55). We show that at times this thoracic muscle complex forms part of flexibly assembled muscle synergies that maintain the postural integrity during moments when the body organizes into gesticulation. As such the expressiveness of gesture lies in part in its biomechanical connection with the voice.

## Supporting information

### Supporting methods

This study has been approved by the Ethics Committee Social Sciences (ECSS) of the Radboud University (reference nr.: 22N.002642). This study has been pre-registered before data collection (see <u>preregistration</u>). This supporting information is part of a computationally reproducible Rmarkdown methods and results pipeline that can be accessed via https://wimpouw.github.io/kineticsvoice/. As such, some pieces of information are repeated here in the supporting information so as to increase the self-contained nature of this computationally reproducible document.

### Experimental design

This study concerned a two-level wrist-weight manipulation (no weight vs. weight), a two-level within-subject vocalization condition (expire vs.vocalize), and a five-level within-subject movement condition (‘no movement’, ‘extension’, ‘flexion’, ‘external rotation’, ‘internal rotation’). With 4 trial repetitions over the experiment, we yield 80 (2 weights x 2 vocalizations x 5 movements x 4 repetitions) trials per participant. Trials were blocked by weight condition and vocalization condition (so that weight and task conditions did not switch constantly from trial to trial). Within blocks all movement conditions were randomized.

### Participants

For the current pre-registered confirmatory experiment, as planned and supported by a power analysis (see preregistration), we collected *N* = (17) participants: 7 female, 10 male, *M* (*SD*) age = 28.50 (6.50), *M* (*SD*) body weight = 72.10kg (10.20), *M* (*SD*) body height = 175.10 cm (8.50), *M*(*SD*) BMI = 23.40 (2.20), *M* (*SD*) triceps skinfold = 19.10mm (4.30).

### Exclusions and deviations from pre-registration

We also performed the experiment with one other participant, but due to an issue with LSL streaming, this dataset could not be synchronized and was lost. Furthermore, due to running over time, one participant had to terminate the study earlier about halfway through. Note further, that in our pre-registration we wanted to admit participants with a BMI lower than 25, but since participants were difficult to recruit we accepted three participants with slightly higher BMIs too (max BMI of the current dataset: 27.10). Participants were all able-bodied and did not have any constraints in performing the task. Finally, as stated in the pre-registration we will only report results on the vocalization trials (and not the expiration-only trials). This means that for this report in total, we have N = 17 participants, with 636 analyzable trials, with balanced conditions: weight = 50.20%, no movement = 19.70%, internal rotation = 20.00 %, external rotation = 20.30%, flexion = 20.10 %, extension = 20.00 %.

### Measurements and equipment

#### Body measurements

To enable future analyses of possible modulating individual-specific body properties we collect some basic information about body properties. Namely, weight, under-arm length, upper-arm length, triceps skinfold, and upper-arm circumference.

#### Experiment protocol

The experiment was coded in Python using functions from PsychoPy. The experiment was controlled via a Brainvision Button Box (Brain Products GmbH, Munich, Germany), which was also streaming its output to the data collection PC unit.

#### Wrist weights

To manipulate the mass set in motion, we apply a wrist weight. We use a TurnTuri sports wrist weight of 1 kg.

#### Video and kinematics

The participants are recorded via a video camera (Logitech StreamCam), sampling at 60 frames per second. We used Mediapipe (56) to track the skeleton and facial movements, which is implemented in Masked-piper which we also use for masking the videos (57). The motion-tracked skeleton, specifically the wrist of the dominant hand, is used to estimate movement initiation, peak speed, and the end of the movement. The motion tracking is however only used for determining movement windows and is not of central concern.

#### Muscle activity (surface ElectroMyography: EMG)

We measured surface EMG using a wired BrainAmp ExG system (Brain Products GmbH, Munich, Germany). Disposable surface electrodes (Kendall 24mm Arbo H124SG) were used, and for each of the four muscle targets we had 3 (active, reference, ground) electrodes (12 electrodes total). The sEMG system sampled at 2500 Hz (for post-processing filters see below).

For an overview of the electrode attachments see Figure S1. We prepare the skin surface for EMG application with a scrub gel (NuPrep) followed by a cotton ball swipe with alcohol (Podior 70%). Active and reference electrodes were attached with a 15mm distance center to center.

We attached electrodes for focal muscles that directly participate in the internal (pectoralis major) and external rotation (infraspinatus) of the humerus. Electrodes were applied for focal muscles ipsilaterally (relative to the dominant hand). We attached electrodes to the muscle belly of the clavicular head of the pectoralis major, with a ground electrode on the clavicle on the opposite side.

We also attached electrodes for postural muscles which will likely anticipate and react to postural perturbations due to upper limb movements. Since these muscles should act in the opposite direction of the postural perturbation of the dominant hand, we applied electrodes contralaterally to the dominant hand. We attach electrodes to the rectus abdominis, with a ground electrode on the iliac crest on the opposite side. We also attached electrodes to the erector spinae muscle group (specifically, the iliocostalis lumborum).

**Figure S1.**
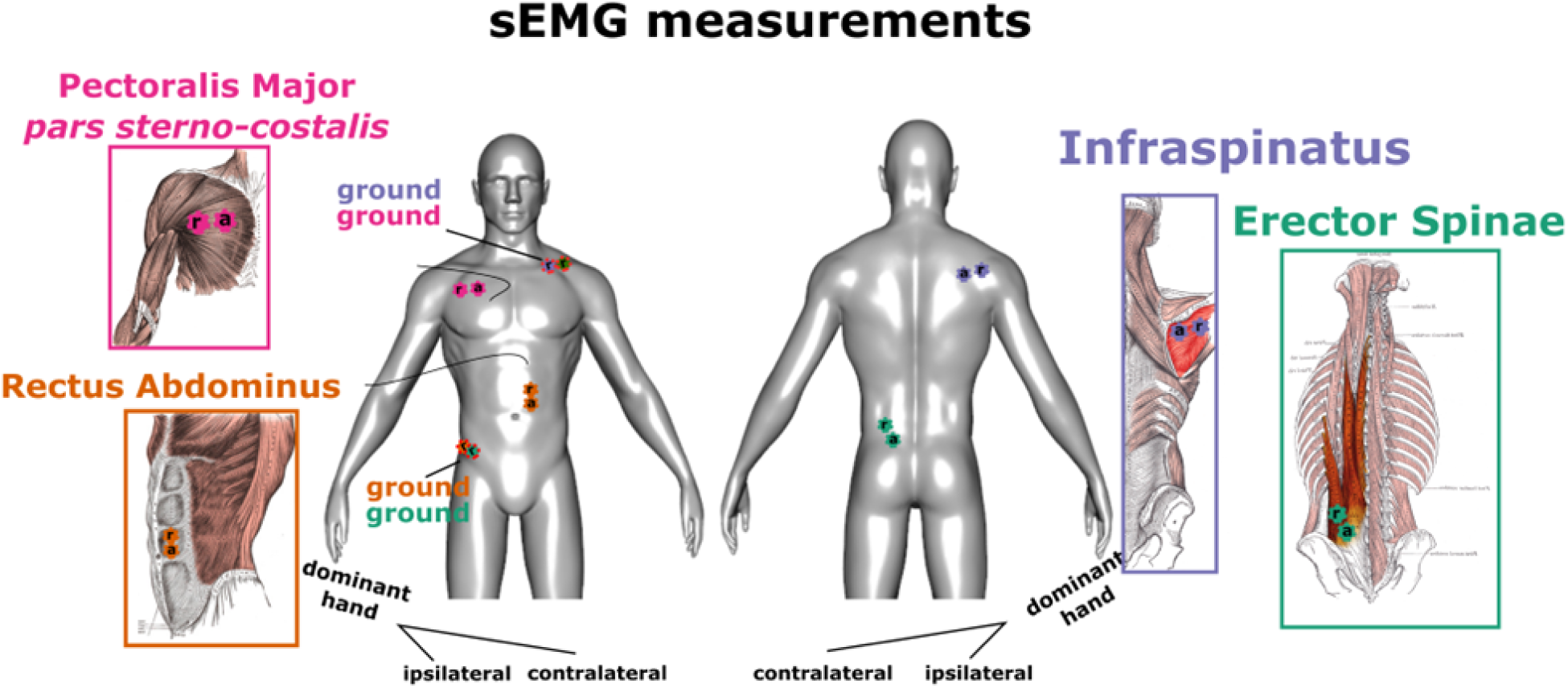
Overview sEMG target muscles. Active (a) reference (r), and ground (g) sEMG electrodes, for each muscle target.

#### Ground reaction measurements

We used an inhouse-built 1m² balance board with vertical pressure sensors. The sensors were derived and remodified from the Wii-Balance board sensors. The sampling rate was 400 Hz. The system was time-locked within millisecond accuracy and has a spatial accuracy of several sub-millimeters. A National Instruments card, USB-62221 performed the A/D conversion and was connected via USB to the PC.

#### Acoustics

To ensure proper acoustic intensity measurements we used a headset microphone; MicroMic C520 (AKG, Inc.) headset condenser cardioid microphone sampling at 16 kHz. The gain levels of the condenser power source were set by the hardware (and could not be changed).

#### Recording setup and synchronization

<u>LabStreamLayer</u> provided a uniform interface for streaming different signals along a network, where a common time stamp for each signal ensures sub-millisecond synchronization. We used a Linux system to record and stream the microphone recordings. Additionally a second PC collected video, and streamed ground reaction forces, and EMG. A data collection PC collected the audio, ground reaction force, and EMG streams and stored the output in XDF format for efficient storing of multiple time-varying signals. The video data was synchronized by streaming the frame number to the LSL recorder, allowing us to match up individual frames with the other signals (even when a frame is dropped).

#### Procedure

Participants were admitted to the study based on exclusion criteria and signed an informed consent. We asked participants to take off their shoes and we proceeded with the body measurements, while instructing the participant about the nature of the study. After body measurements, we applied the surface EMG. We prepared the muscle site with rubbing gel and alcohol, and active/reference electrodes were placed with a distance of 15 mm from each other’s center. See Figure S1 for the sEMG electrode locations. The procedures up to the start of the experiment take about 20 minutes or less in total.

Upon the start of the experiment participants took a standing position on the force platform. The experiment commenced with calibration and practice trials. First, 10 seconds of silent breathing without body movements were recorded. Then participants were asked to take a maximum inspiration followed by a maximum expiration, so as to measure signal conditions under respiratory boundary conditions. Then, for the practice trials, each movement was practiced with expiring and vocalization while performing the movement conditions, and the participant is introduced to wearing the wrist weight of 1 kg. After practice trials, participants performed 80 blocked trials.

For each (practice) trial participants were closely guided by the information on the monitor. Firstly, participants were shown the movement to be performed for the trial, and were prompted by the experimenter to get ready to repeat the movement. No detailed instructions were given about how to move, other than showing the movements in animations, showing a person-masked skeleton performing the movement. Then participants were instructed to adopt the start position of the movement, which is a 90 degree elbow flexion, with either an externally rotated humerus (start position for internal rotation), or a non-rotated humerus with the wrist in front of the body (rest position for the other movement conditions). For the no movement condition, participants were asked to rest their arms alongside their body. Upon trial start, participants inhaled deeply with a timer counting down from 4 seconds. Then, participants vocalized or expired depending on condition assignment, with a screen appearing after 3 seconds to perform the movement. Visual guidance was provided to remind the participant of the end location, where a static image was shown of the movement end position. After an additional 4 seconds the trial ended. The 4 seconds allowed more than enough time to perform the movement and stabilize vocalization after the perturbation. In the no movement condition, a prompt was given to maintain one’s posture with an image of the rest position (arms alongside the body).

### Preprocessing of the data streams

#### EMG

To reduce heart rate artifacts we applied a common method [@drakeEliminationElectrocardiogramContamination2006] of high-pass filtering the signal at 30 Hz using a zero-phase 4th order Butterworth filter. We then full-wave rectified the EMG signal and applied a zero-phase low-pass 4th-order Butterworth filter at 20 Hz. When filtering any signal we pad the signals to avoid edge effects. We normalized the EMG signals within participants before submitting them to analyses.

**Figure S2.**
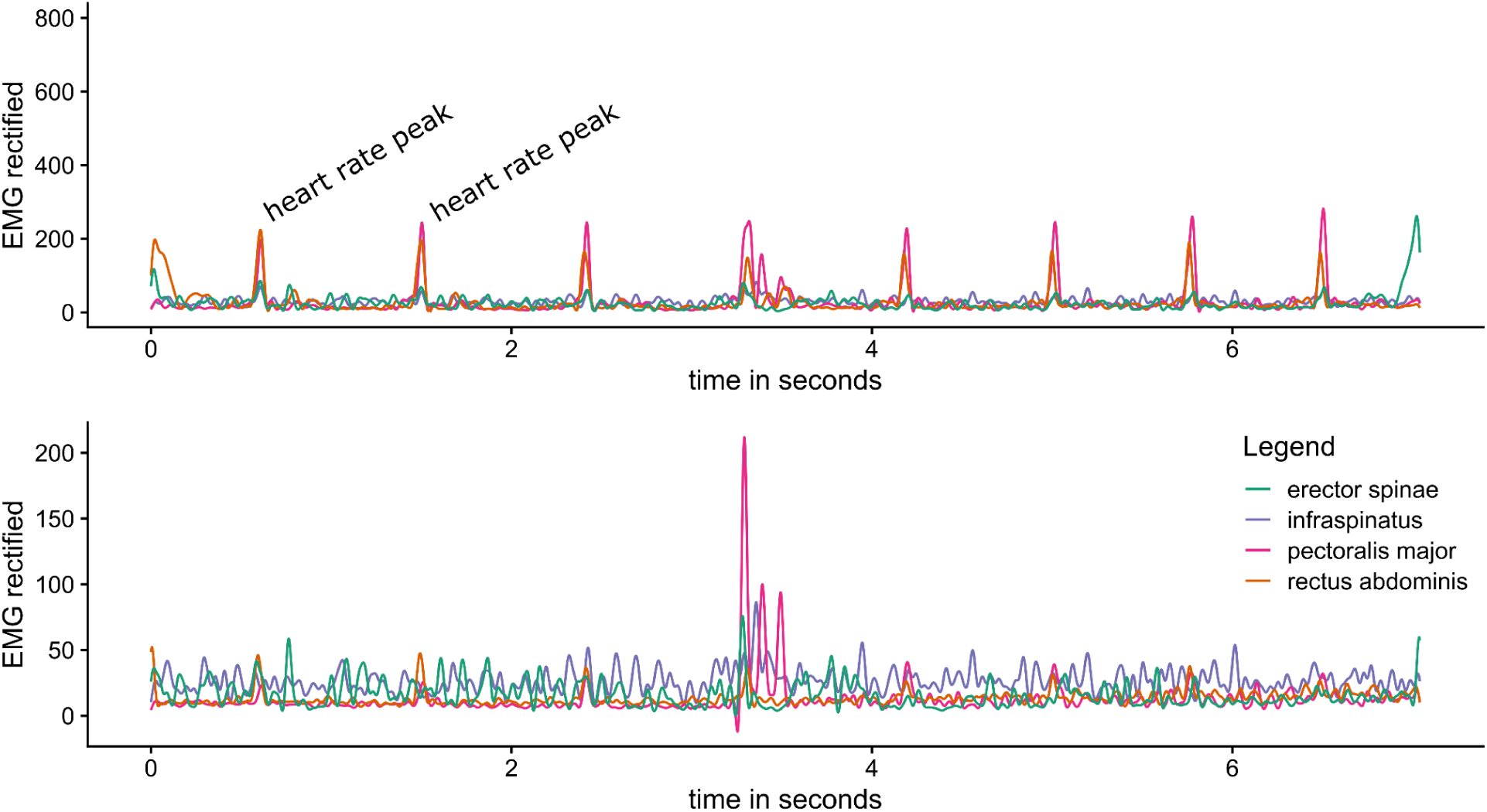
Example of smoothing settings for EMG signals. The upper panel shows the raw rectified high-pass filtered EMG, and the lower panel shows the low-pass filtered data to reduce artifacts of heart rate. This example shows an internal rotation trial, where we successfully retrieve the peak in the pectoralis major that internally rotates the arm.

#### Ground reaction forces

We upsampled the balanceboard from 400 Hz to 2,500 Hz. We then applied a zero-phase low-pass 2nd order Butterworth filter at 20 Hz to the padded signals. These signals were used to calculate the key measure for postural perturbation we computed the change in 2D magnitude (L2 norm for x and y, and combined) in center of pressure. We refer to this as the change in the center of pressure (COPc), or the anterior-posterior COPc (front-back sway), or the medial-lateral COPc (left-right sway).

#### Acoustics

For acoustics, we extracted the smoothed amplitude envelope (hereafter envelope). For the envelope we applied a Hilbert transform to the waveform signal, took the complex modulus to create a 1D time series, which was then resampled at 2,500 Hz, and smoothed with a Hann filter based on a Hanning Window of 12 Hz. We normalize the amplitude envelope signals within participants before submitting them to analyses.

#### Data aggregation

All signals were sampled at, or upsampled to, 2500 Hz. Then we aggregated the data by aligning time series in a combined by-trial format to increase ease of combined analyses. We linearly interpolated signals when sample times did not align perfectly.

#### Data sharing & Privacy

Video data is deidentified using the masked-piper tool to mask faces and bodies while maintaining kinematic information (31).

### Overview data analyses

Note that there is a general decline in the amplitude of the vocalization during a trial (see Fig. S3, top panel). This is to be expected, as the subglottal pressure falls when the lungs deflate. To quantify deviations from stable vocalizations, we therefore detrended the amplitude envelope time series, to assess positive or negative peaks relative to this trend line. For the envelope, muscle activity, and the change in center of pressure, we measure the global maxima happening within the analyses window (i.e., within a trial we take a local maximum occurring between movement onset and offset). We analyzed positive and negative peaks within the movement window separately.

**Figure S3:**
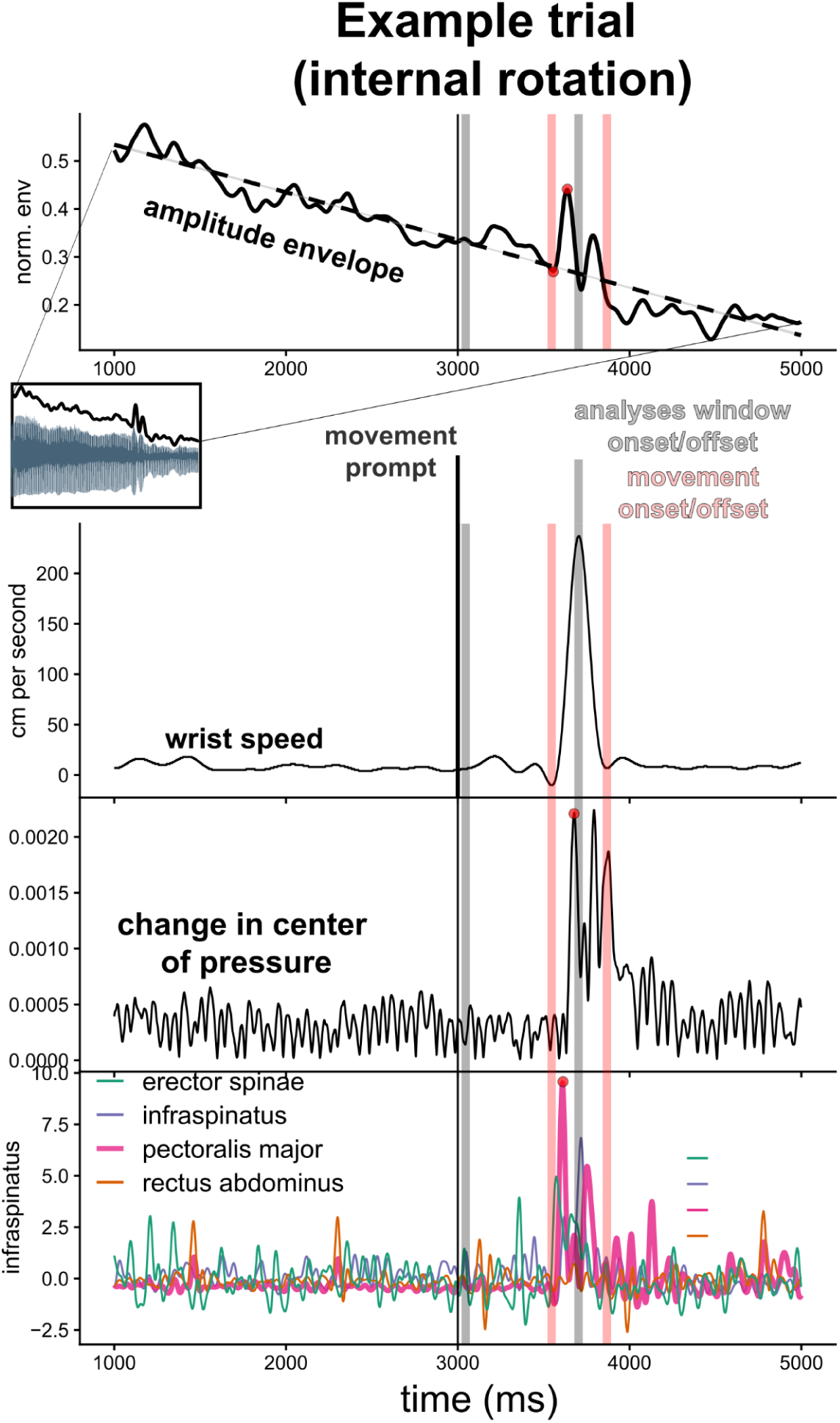
Combined time series. One example trial and the associated signals are shown, for the internal rotation movement condition. At time = 0 s, the prompt is given to the participant to vocalize. We determine a detrending line using linear regression for the 1 to 5 seconds after the vocalization prompt. Note, that at 3 s (3000 ms), there is a movement prompt. However, we determine our window where we assess peaks in signals at 500 ms before and after the movement onset/offset (using peakfinding function on the 2D speed time series of the wrist). In these trials, the analysis window is given in gray dashed bars, which is 500 ms after and before movement onset/offset.

## Supporting results

### Descriptive results and statistical manipulation checking

Before moving to our main pre-registered hypothesis testing, we first provide an overview of the data and statistical manipulation checks. This includes the timing of the movements, the magnitude of the muscle, postural, and kinematic peaks per movement and weight condition, as well as the effect of the vocal condition on the magnitude of those peaks.

### Muscle, postural, and kinematic changes for different movement and weight conditions

Figure S4 shows an overview of the muscle activity patterns for each movement condition, split across weight conditions. Table S1 provides the numerical information. All in all, these descriptive results pattern in sensible ways. There is generally more (postural) muscle, postural, and kinematic activity in movement conditions. Focal muscles powering internal (pectoralis major) or external (infraspinatus) rotation are peaking in activity during these movement conditions. Especially for flexion and extension (rather than internal/external rotation) we obtain anterior-posterior change in the center of pressure. Generally, there is a higher muscle, postural activity when wearing weights, while weights generally reduce peak speed and acceleration. Below we go into more detail, for the statistical confirmation of these findings.

**Figure S4.**
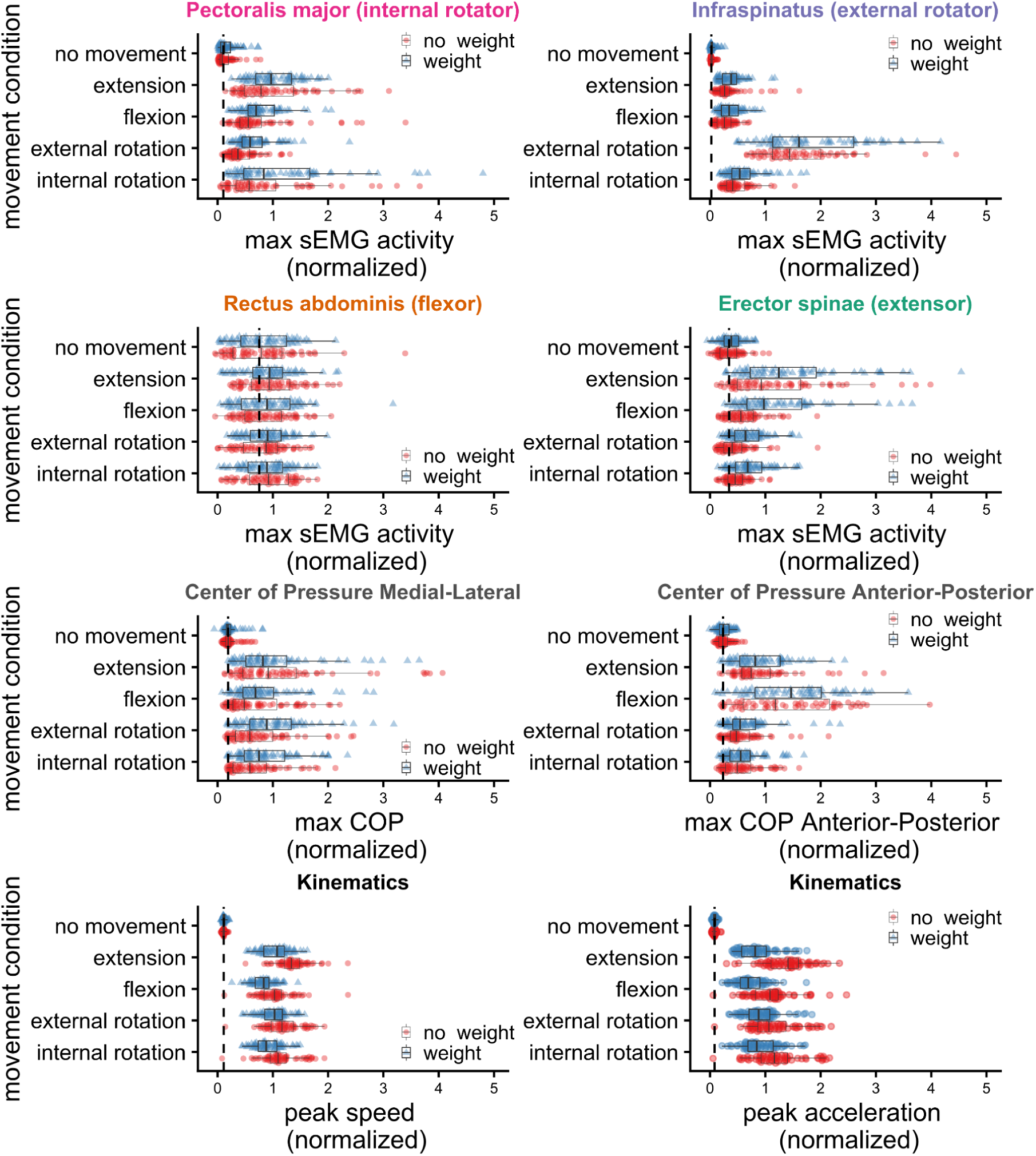
Muscle, postural, and kinematic changes for different movement and weight conditions.

**Table S1.** Numerical information (z-scaled values) related to Figure S4 (or Figure 2 in the main report). COP_ML = Change in center of pressure medial-lateral, COP_AP = COPc anterior-posterior. Values are presented as: Mean (SD) [95% CI lower, upper].

| movement condition<br>weight condition | Pectoralis | Infraspinatus | Rectus | Erector | COPc<br>medial-lateral | COPc<br>anterior-posteri<br>or | Peak Speed | Peak<br>Acceleration |
| --- | --- | --- | --- | --- | --- | --- | --- | --- |
| no movement<br>no weight | 0.15 (+-0.15)<br>[0.11 - 0.18] | 0.03 (+-0.03)<br>[0.03 - 0.04] | 0.81 (+-0.63)<br>[0.65 - 0.97] | 0.35 (+-0.22)<br>[0.29 - 0.41] | 0.22 (+-0.14)<br>[0.19 - 0.26] | 0.25 (+-0.12)<br>[0.22 - 0.29] | 0.11 (+-0.03)<br>[0.10 - 0.12] | 0.08 (+-0.03)<br>[0.07 - 0.09] |
| **no movement**<br>**weight** | **0.18 (+-0.17)**<br>**[0.13 - 0.22]** | **0.04 (+-0.05)**<br>**[0.03 - 0.06]** | **0.81 (+-0.50)**<br>**[0.68 - 0.93]** | **0.39 (+-0.20)**<br>**[0.34 - 0.45]** | **0.23 (+-0.15)**<br>**[0.19 - 0.27]** | **0.26 (+-0.12)**<br>**[0.22 - 0.29]** | **0.12 (+-0.03)**<br>**[0.11 - 0.13]** | **0.08 (+-0.03)**<br>**[0.08 - 0.09]** |
| extension<br>no weight | 0.99 (+-0.67)<br>[0.82 - 1.16] | 0.32 (+-0.28)<br>[0.25 - 0.39] | 0.98 (+-0.50)<br>[0.85 - 1.10] | 1.17 (+-0.88)<br>[0.95 - 1.40] | 1.17 (+-0.98)<br>[0.92 - 1.42] | 0.92 (+-0.58)<br>[0.77 - 1.06] | 1.36 (+-0.28)<br>[1.29 - 1.43] | 1.38 (+-0.40)<br>[1.28 - 1.48] |
| **extension**<br>**weight** | **1.03 (+-0.43)**<br>**[0.92 - 1.14]** | **0.38 (+-0.22)**<br>**[0.33 - 0.44]** | **0.91 (+-0.48)**<br>**[0.79 - 1.03]** | **1.46 (+-0.94)**<br>**[1.23 - 1.70]** | **1.02 (+-0.78)**<br>**[0.82 - 1.22]** | **0.90 (+-0.49)**<br>**[0.78 - 1.03]** | **1.04 (+-0.26)**<br>**[0.98 - 1.10]** | **0.84 (+-0.32)**<br>**[0.76 - 0.92]** |
| flexion<br>no weight | 0.75 (+-0.64)<br>[0.59 - 0.91] | 0.27 (+-0.16)<br>[0.23 - 0.31] | 0.85 (+-0.46)<br>[0.73 - 0.96] | 0.62 (+-0.35)<br>[0.53 - 0.70] | 0.71 (+-0.55)<br>[0.57 - 0.84] | 1.40 (+-0.88)<br>[1.18 - 1.62] | 1.04 (+-0.32)<br>[0.96 - 1.12] | 1.09 (+-0.40)<br>[0.99 - 1.19] |
| **flexion**<br>**weight** | **0.79 (+-0.38)**<br>**[0.69 - 0.88]** | **0.37 (+-0.21)**<br>**[0.32 - 0.42]** | **0.91 (+-0.56)**<br>**[0.77 - 1.05]** | **1.23 (+-0.77)**<br>**[1.03 - 1.42]** | **0.85 (+-0.58)**<br>**[0.70 - 0.99]** | **1.48 (+-0.81)**<br>**[1.28 - 1.69]** | **0.83 (+-0.21)**<br>**[0.78 - 0.88]** | **0.73 (+-0.27)**<br>**[0.66 - 0.80]** |
| external rotation<br>no weight | 0.44 (+-0.28)<br>[0.37 - 0.51] | 1.59 (+-0.76)<br>[1.40 - 1.78] | 0.81 (+-0.43)<br>[0.70 - 0.92] | 0.46 (+-0.32)<br>[0.38 - 0.54] | 0.75 (+-0.56)<br>[0.61 - 0.89] | 0.57 (+-0.36)<br>[0.48 - 0.66] | 1.19 (+-0.32)<br>[1.11 - 1.27] | 1.16 (+-0.42)<br>[1.06 - 1.27] |
| **external rotation**<br>**weight** | **0.68 (+-0.38)**<br>**[0.59 - 0.78]** | **1.98 (+-1.19)**<br>**[1.68 - 2.27]** | **0.90 (+-0.40)**<br>**[0.80 - 1.00]** | **0.69 (+-0.33)**<br>**[0.61 - 0.77]** | **1.01 (+-0.63)**<br>**[0.85 - 1.16]** | **0.66 (+-0.42)**<br>**[0.55 - 0.76]** | **1.01 (+-0.25)**<br>**[0.95 - 1.08]** | **0.92 (+-0.32)**<br>**[0.84 - 1.00]** |
| internal rotation<br>no weight | 0.93 (+-0.99)<br>[0.68 - 1.19] | 0.48 (+-0.26)<br>[0.41 - 0.54] | 0.89 (+-0.44)<br>[0.78 - 1.00] | 0.48 (+-0.21)<br>[0.43 - 0.53] | 0.67 (+-0.47)<br>[0.55 - 0.79] | 0.57 (+-0.35)<br>[0.48 - 0.66] | 1.14 (+-0.34)<br>[1.05 - 1.23] | 1.22 (+-0.46)<br>[1.11 - 1.34] |
| **internal rotation**<br>**weight** | **1.24 (+-1.11)**<br>**[0.96 - 1.52]** | **0.61 (+-0.34)**<br>**[0.53 - 0.70]** | **0.88 (+-0.42)**<br>**[0.78 - 0.99]** | **0.74 (+-0.36)**<br>**[0.65 - 0.83]** | **0.88 (+-0.53)**<br>**[0.75 - 1.01]** | **0.61 (+-0.35)**<br>**[0.52 - 0.70]** | **0.91 (+-0.25)**<br>**[0.85 - 0.98]** | **0.91 (+-0.33)**<br>**[0.83 - 0.99]** |

### Statistical tests for manipulation checking of movement vs. no movement condition on muscle and postural measurements

First, we report whether our manipulation of movement vs. no movement (binary comparison) indeed increases postural and muscle activity. As expected, the increased activity of muscles, except the rectus abdominis, is confirmed by comparing mixed regression models (participant as random intercept) predicting the magnitude of the peak muscle activity with a movement vs. no movement predictor included, versus a model predicting the overall mean.

**Table S2.** Muscle and postural activity predicted by presence of movement versus no movement.

| Measure | Chi2 | df | pvalue |
| --- | --- | --- | --- |
| movement vs. no movement (pectoralis) | 261.2025 | 1 | 0.0000 |
| movement vs. no movement (infraspinatus) | 199.4645 | 1 | 0.0000 |
| movement vs. no movement (rectus abdominis) | 3.7817 | 1 | 0.0518 |
| movement vs. no movement (erector spinae) | 113.2997 | 1 | 0.0000 |
| movement vs. no movement (COPc) | 360.9009 | 1 | 0.0000 |

### Statistical tests for manipulation checking of differences between movement-only conditions on muscle, postural, and kinematic measurements

We assessed whether there is an effect of weight in movement-only conditions over and above a model that includes a movement predictor. We find that weight indeed affects all measured muscle activity except the rectus abdominis. Weight also affects kinematics, confirming lower speeds and accelerations when moving with a weight (Table S1, Figure S4). However, we did not find statistically reliable effects of weight on the change in the center of pressure.

**Table S3.** Muscle activity predicted by presence of weight in the movement conditions over and above an effect of movement condition.

| Muscle | Chi2 | df | pvalue |
| --- | --- | --- | --- |
| Weight vs. no weight (pectoralis major) | 34.7011 | 1 | 0.0000 |
| Weight vs. no weight (infraspinatus) | 15.6817 | 1 | 0.0001 |
| Weight vs. no weight (rectus abdominis) | 0.0838 | 1 | 0.7721 |
| Weight vs. no weight (erector spinae) | 79.1906 | 1 | 0.0000 |
| Weight vs. no weight (wrist speed) | 204.4295 | 1 | 0.0000 |
| Weight vs. no weight (wrist acceleration) | 257.7740 | 1 | 0.0000 |
| Weight vs. no weight (COP) | 1.2612 | 1 | 0.2614 |

### Statistical tests for manipulation checking of differences between movement-only conditions on muscle, postural, and kinematic measurements

We performed the same type of manipulation check to assess whether there are any differences in posture (anterior posterior, medial-lateral), muscle activity, and kinematics (speed and acceleration), between the different types of movements (i.e., the movement-only conditions). We only test overall effects of movement conditions, versus a model predicting the overall mean (but also see table S1 and Figure S4 for detailed information). Different types of movements lead to different peak muscle activity (except for the rectus abdominis), as well as leading to different peak change of center of pressure and kinematics (speed and acceleration).

**Table S4.**
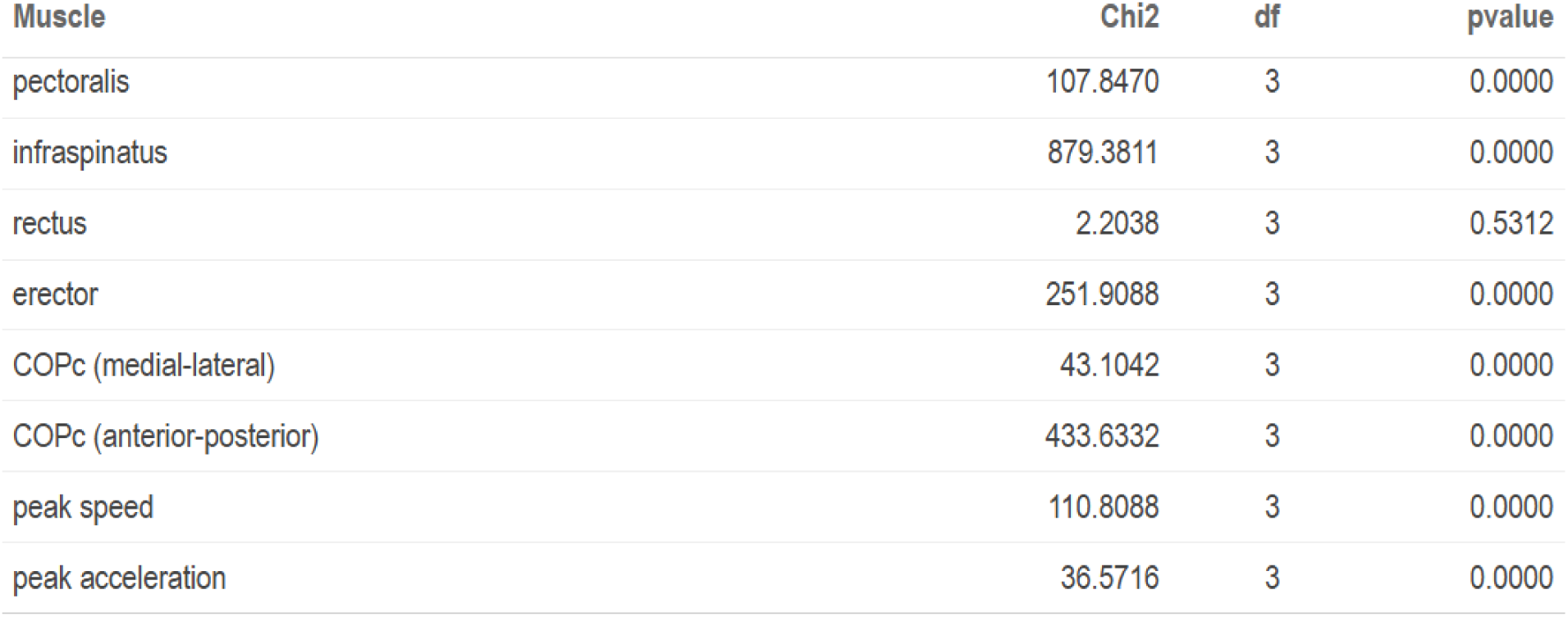
Muscle and postural activity predicted by different types of movements versus a base model.

| Muscle | Chi2 | df | pvalue |
| --- | --- | --- | --- |
| pectoralis | 107.8470 | 3 | 0.0000 |
| infraspinatus | 879.3811 | 3 | 0.0000 |
| rectus | 2.2038 | 3 | 0.5312 |
| erector | 251.9088 | 3 | 0.0000 |
| COPc (medial-lateral) | 43.1042 | 3 | 0.0000 |
| COPc (anterior-posterior) | 433.6332 | 3 | 0.0000 |
| peak speed | 110.8088 | 3 | 0.0000 |
| peak acceleration | 36.5716 | 3 | 0.0000 |

## Main Confirmatory Analyses

### Research Question 1: Do movement and weight conditions have (different) effects on vocalization amplitude?

From inspecting the summarized trajectories, and the descriptive exploratory analyses above, we obtain that the internal rotation of the arm seems to be preceded by a negative peak in vocalization around the onset of the movement, which is followed by a positive peak in the voice amplitude. Furthermore, we observe that all other movements primarily have a positive peak in the amplitude of the voice. A straightforward test of whether the voice amplitude envelope has positive or negative peaks is to assess differences in peaks in the vocalization conditions per movement (and weight) condition.

**Figure S5:**
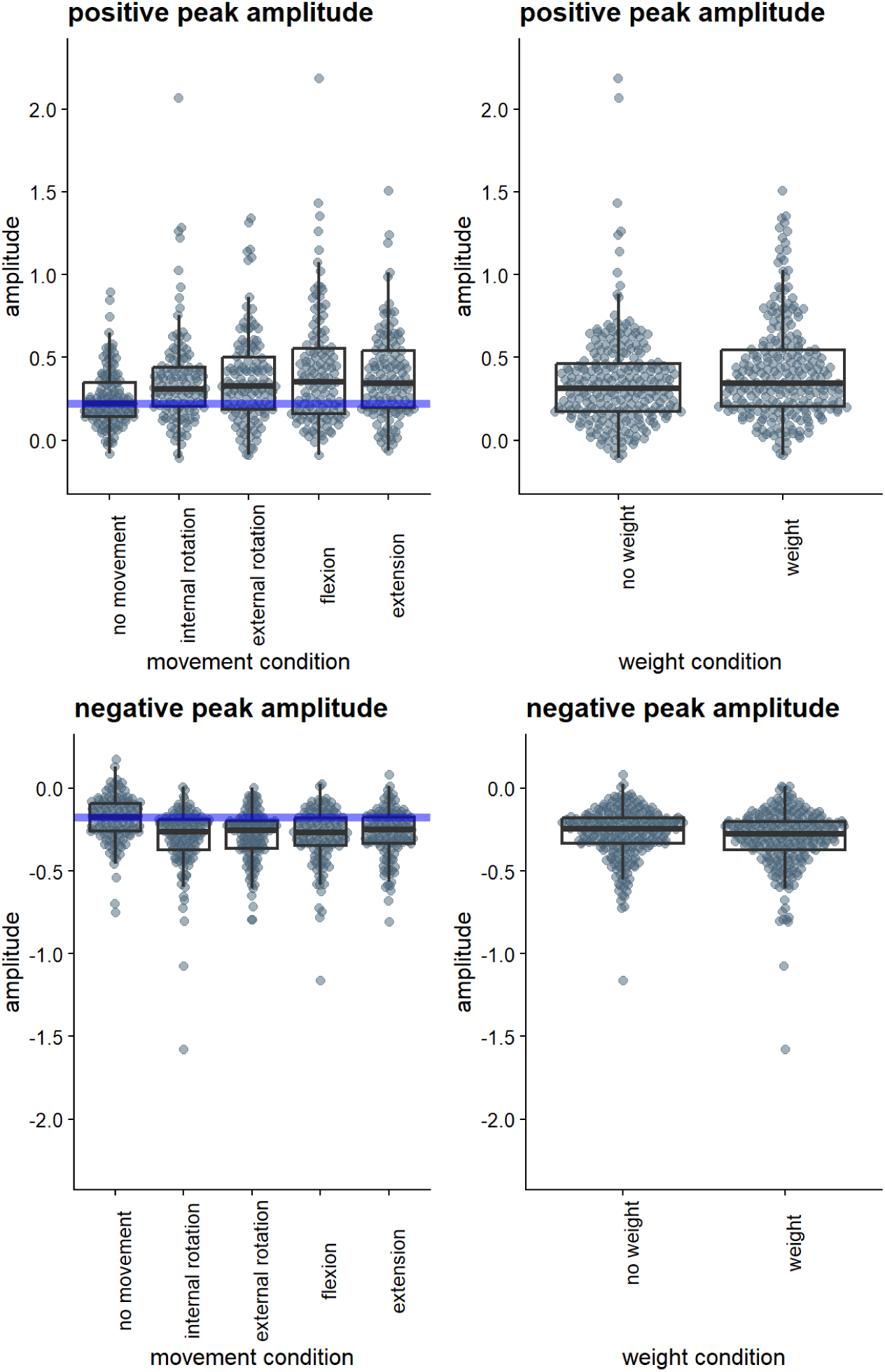
Effects of movement condition of positive and negative peaks in the voice amplitude. Effects of movement condition of positive and negative peaks in the voice amplitude. The upper part shows the positive peaks in the amplitude envelope during the different movement and weight conditions. The lower part shows the negative peaks (hence the negative values; note, in the modeling we will absolutize these values). It is clear that relative to vocalization of no movement, there are especially positive, but also more negative peaks in the amplitude envelope for the different movement conditions.

We first modeled with a mixed linear regression the variation in *positive* peaks in the amplitude envelope (using R-package lme4), with participant as random intercept (for more complex random slope models did not converge). A model with weight and movement conditions explained more variance than a base model predicting the overall mean, Change in χ2 (5) = 36.06, *p* < .001). he model coefficients are given in Table S5. It can be seen that there is a positive but not statistically reliable effect of wrist weight in this sample. Further, all movements (extension, flexion, internal rotation, external rotation) lead to statistically reliable increases in positive peaks in the amplitude envelope relative to the no movement condition (with flexion and external rotation leading to more extreme effects).

**Table S5.** Effects of weight and movement condition on magnitude positive peaks in amplitude envelope.

|  | Estimate | SE | df | t-value | p-value |
| --- | --- | --- | --- | --- | --- |
| Intercept (no movement/no weight) | 0.233 | 0.037 | 42.600 | 6.367 | 0.000 |
| vs. weight | 0.058 | 0.021 | 603.791 | 2.819 | 0.005 |
| vs. internal rotation | 0.104 | 0.033 | 602.728 | 3.202 | 0.001 |
| vs. external rotation | 0.116 | 0.032 | 602.740 | 3.587 | 0.000 |
| vs. flexion | 0.164 | 0.033 | 602.778 | 5.054 | 0.000 |
| vs. extension | 0.130 | 0.032 | 602.717 | 4.008 | 0.000 |

Note that in the previous model we assess the role of weight vs. no weight. But during the no movement condition the weight is not affecting the participant. Thus, a better estimate of the effect of wrist weight is to assess weight vs. no weight conditions for only the movement conditions. Below it shows indeed that this comparison further increases the effect strength as compared to a model with the no movement condition included.

**Table S6:** Effects of weight on magnitude of positive peaks in amplitude envelope (no movement condition filtered out).

|  | Estimate | SE | df | t-value | p-value |
| --- | --- | --- | --- | --- | --- |
| Intercept (no weight) | 0.361 | 0.035 | 20.173 | 10.303 | 0.000 |
| vs. weight | 0.062 | 0.024 | 484.657 | 2.565 | 0.011 |

Secondly, we modeled in a similar way the negative peaks in the amplitude envelope, and found that a model with weight and movement conditions explained more variance than a base model predicting the overall mean, Change in χ2 (5) = 48.47, *p* < .001). Model coefficients are shown in Table S7. We find that some movement conditions (flexion, internal rotation, and especially external rotation) had higher magnitude negative peaks relative to no movement. No reliable effect of weight condition was found, nor did the extension movement lead to negative peaks relative to the no movement condition. Note, in our models, we have absolutized the values of negative peaks, such that positive effects of some condition means higher magnitude negative peaks (i.e., more negative peaks).

**Table S7:**
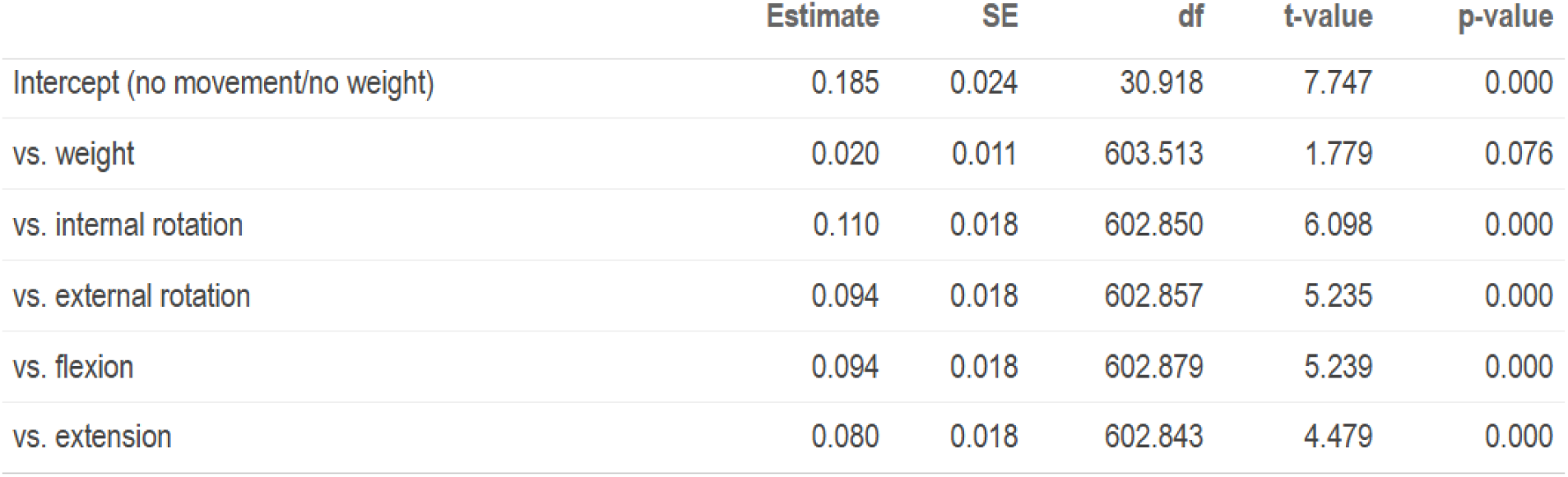
Effects of weight and movement condition on magnitude negative peaks in amplitude envelope.

|  | Estimate | SE | df | t-value | p-value |
| --- | --- | --- | --- | --- | --- |
| Intercept (no movement/no weight) | 0.185 | 0.024 | 30.918 | 7.747 | 0.000 |
| vs. weight | 0.020 | 0.011 | 603.513 | 1.779 | 0.076 |
| vs. internal rotation | 0.110 | 0.018 | 602.850 | 6.098 | 0.000 |
| vs. external rotation | 0.094 | 0.018 | 602.857 | 5.235 | 0.000 |
| vs. flexion | 0.094 | 0.018 | 602.879 | 5.239 | 0.000 |
| vs. extension | 0.080 | 0.018 | 602.843 | 4.479 | 0.000 |

**Table S8:** Effects of weight on magnitude of negative peaks in amplitude envelope (no movement condition filtered out).

|  | Estimate | SE | df | t-value | p-value |
| --- | --- | --- | --- | --- | --- |
| Intercept (no weight) | -0.274 | 0.024 | 18.498 | -11.607 | 0.000 |
| vs. weight | -0.029 | 0.013 | 484.463 | -2.259 | 0.024 |

We further performed an exploratory post-hoc analysis to see if the different movement conditions themselves were reliably different from each other in terms of affecting the voice. We performed Tukey-corrected post-hoc analyses on the linear mixed regression models using R-package ’emmeans’ (58). As can be seen, there are no differences between movement-only conditions in terms of the degree they affect the magnitude of positive or negative peaks in the amplitude envelope of the vocalization.

**Table S9.**
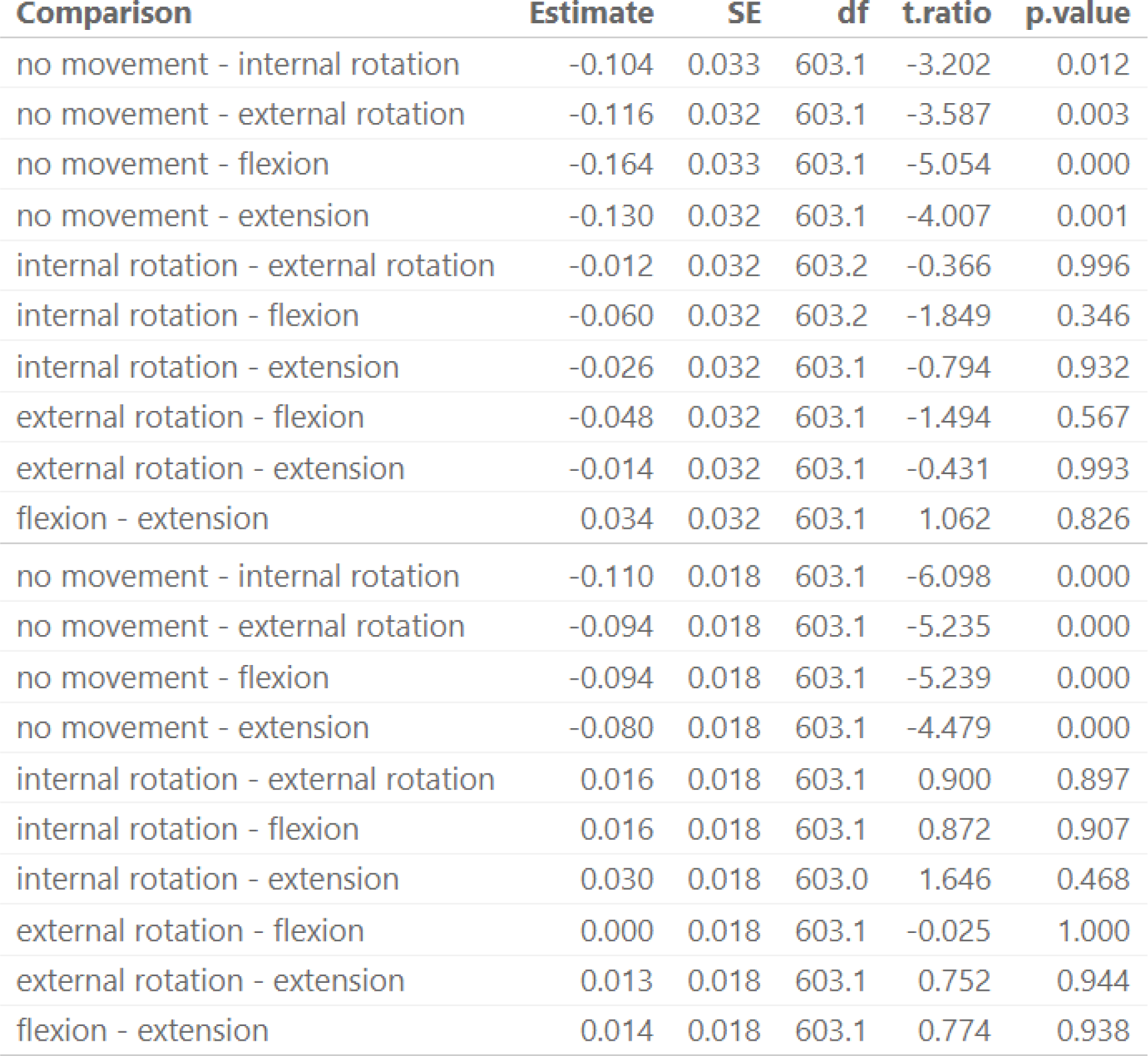
Pairwise Comparisons of Movement Conditions (Positive Peaks)

| Comparison | Estimate | SE | df | t.ratio | p.value |
| --- | --- | --- | --- | --- | --- |
| no movement - internal rotation | -0.104 | 0.033 | 603.1 | -3.202 | 0.012 |
| no movement - external rotation | -0.116 | 0.032 | 603.1 | -3.587 | 0.003 |
| no movement - flexion | -0.164 | 0.033 | 603.1 | -5.054 | 0.000 |
| no movement - extension | -0.130 | 0.032 | 603.1 | -4.007 | 0.001 |
| internal rotation - external rotation | -0.012 | 0.032 | 603.2 | -0.366 | 0.996 |
| internal rotation - flexion | -0.060 | 0.032 | 603.2 | -1.849 | 0.346 |
| internal rotation - extension | -0.026 | 0.032 | 603.1 | -0.794 | 0.932 |
| external rotation - flexion | -0.048 | 0.032 | 603.1 | -1.494 | 0.567 |
| external rotation - extension | -0.014 | 0.032 | 603.1 | -0.431 | 0.993 |
| flexion - extension | 0.034 | 0.032 | 603.1 | 1.062 | 0.826 |

**Table S9.** Pairwise Comparisons of Movement Conditions (Positive Peaks)
| Comparison | Estimate | SE | df | t.ratio | p.value |
| --- | --- | --- | --- | --- | --- |
| no movement - internal rotation | -0.110 | 0.018 | 603.1 | -6.098 | 0.000 |
| no movement - external rotation | -0.094 | 0.018 | 603.1 | -5.235 | 0.000 |
| no movement - flexion | -0.094 | 0.018 | 603.1 | -5.239 | 0.000 |
| no movement - extension | -0.080 | 0.018 | 603.1 | -4.479 | 0.000 |
| internal rotation - external rotation | 0.016 | 0.018 | 603.1 | 0.900 | 0.897 |
| internal rotation - flexion | 0.016 | 0.018 | 603.1 | 0.872 | 0.907 |
| internal rotation - extension | 0.030 | 0.018 | 603.0 | 1.646 | 0.468 |
| external rotation - flexion | 0.000 | 0.018 | 603.1 | -0.025 | 1.000 |
| external rotation - extension | 0.013 | 0.018 | 603.1 | 0.752 | 0.944 |
| flexion - extension | 0.014 | 0.018 | 603.1 | 0.774 | 0.938 |

### Research Question 2: Do movement and weight conditions have (different) effects on vocalization amplitude?

Since each movement and weight condition is designed to recruit different muscle activations, we can also directly relate muscle activity peaks with the positive and negative peaks in the amplitude envelope. We use a similar linear mixed regression approach to model variance in vocal amplitude peaks with peaks in muscle activity for the different muscles measured. Since there are correlations between these muscle activities (see Tables S13, S14, S15, S16), that could affect effect estimation in linear mixed regression, we first assessed the Variance Inflation Factors (VIF) between the muscle activity peaks. This yielded a maximum VIF value of 1.28. Since this is considered a low value (VIF > 5 is generally considered problematic), we can combine the different muscle activity measurements in one model to predict amplitude envelope peaks. Figure S6 shows the graphical results of these relationships

**Figure S6.**
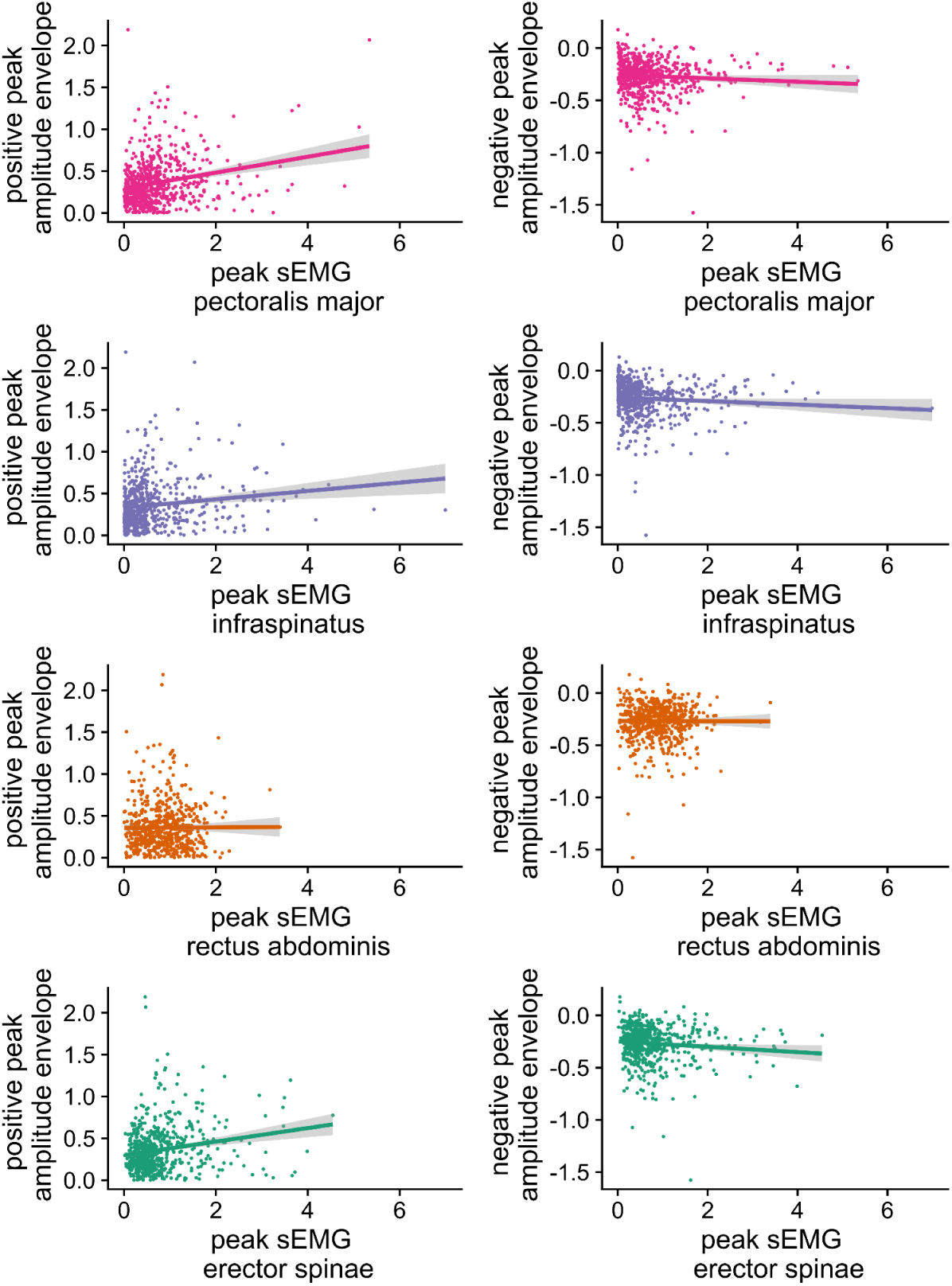
Relations between peak muscle activity and positive peaks in the amplitude envelope.

In a model with participant as random intercept the different peak muscle activities explained more variance than a base model predicting the overall mean, Change in χ2 = 59.08 (4), *p* < .001). We also tried to fit a more complex model with random slopes for participants, but such a model did not converge. The model coefficients are given in Table S10. Further, Table S10 shows that peak EMG activity in all the muscles (except for the rectus abdominis) leads to statistically reliable increases in positive peaks in the amplitude envelope (with stronger effects).

**Table S10.** Linear mixed regression model assessing the relation of peak muscle activity with the positive peak in the amplitude envelope.

|  | Estimate | SE | df | t-value | p-value |
| --- | --- | --- | --- | --- | --- |
| Intercept | 0.240 | 0.037 | 54.507 | 6.436 | 0.000 |
| erector spinae | 0.016 | 0.004 | 617.719 | 4.208 | 0.000 |
| infraspinatus | 0.006 | 0.002 | 612.274 | 2.822 | 0.005 |
| pectoralis major | 0.013 | 0.003 | 617.427 | 4.809 | 0.000 |
| rectus abdominis | -0.003 | 0.007 | 612.555 | -0.373 | 0.710 |

We similarly modeled the variance of the magnitude in negative peaks in the amplitude envelope with muscle activity peaks (see right side of Figure S7), and we similarly observed that such a model performed better than a base model predicting the overall mean, Change in χ2 = 16.48 (4.00), *p* = 0.002). The model coefficients are given in Table S11. Further, as shown in Table S8 for the infraspinatus and the pectoralis major, increases in peak EMG activity lead to statistically reliable increases in the magnitude of negative peaks in the amplitude envelope (with stronger effects), while the effects of erector spinae and the rectus abdominis do not reach statistical reliability.

**Table S11:** Linear mixed regression model assessing the relation of peak muscle activity with the negative peak in the amplitude envelope. The negative peaks in the amplitude envelope are absolutized.

|  | Estimate | SE | df | t-value | p-value |
| --- | --- | --- | --- | --- | --- |
| Intercept | 0.227 | 0.026 | 36.136 | 8.826 | 0.000 |
| erector spinae | 0.004 | 0.002 | 613.139 | 1.962 | 0.050 |
| infraspinae | 0.003 | 0.001 | 608.855 | 2.330 | 0.020 |
| pectoralis major | 0.003 | 0.002 | 612.837 | 2.143 | 0.033 |
| rectus abdominis | 0.001 | 0.004 | 619.957 | 0.354 | 0.723 |

### Research Question 3: Is postural muscle activity related to postural stability (change in center of mass)?

Finally, we will assess which muscle activity can be related to changes in the center of pressure, which would directly confirm that gesture-speech biomechanics relate to postural stability (12). Figure S7 shows the graphical results. We similarly performed a linear mixed regression (with participant as random intercept) with a model containing peak EMG activity for each muscle which was regressed on the peak in the change in the center of pressure (COPc). We obtained that a base model predicting the overall mean of COPc was outperformed relative to said model, Change in χ2 (4.00) = 180.01, *p* < .001. Table S12 provides the coefficient information. We find that only the postural muscles (rectus abdominis, erector spinae) indeed reliably predict the magnitude of changes in the center of pressure, while the pectoralis major and the infraspinatus do not reliably relate to the changes in ground reaction forces.

**Figure S7.**
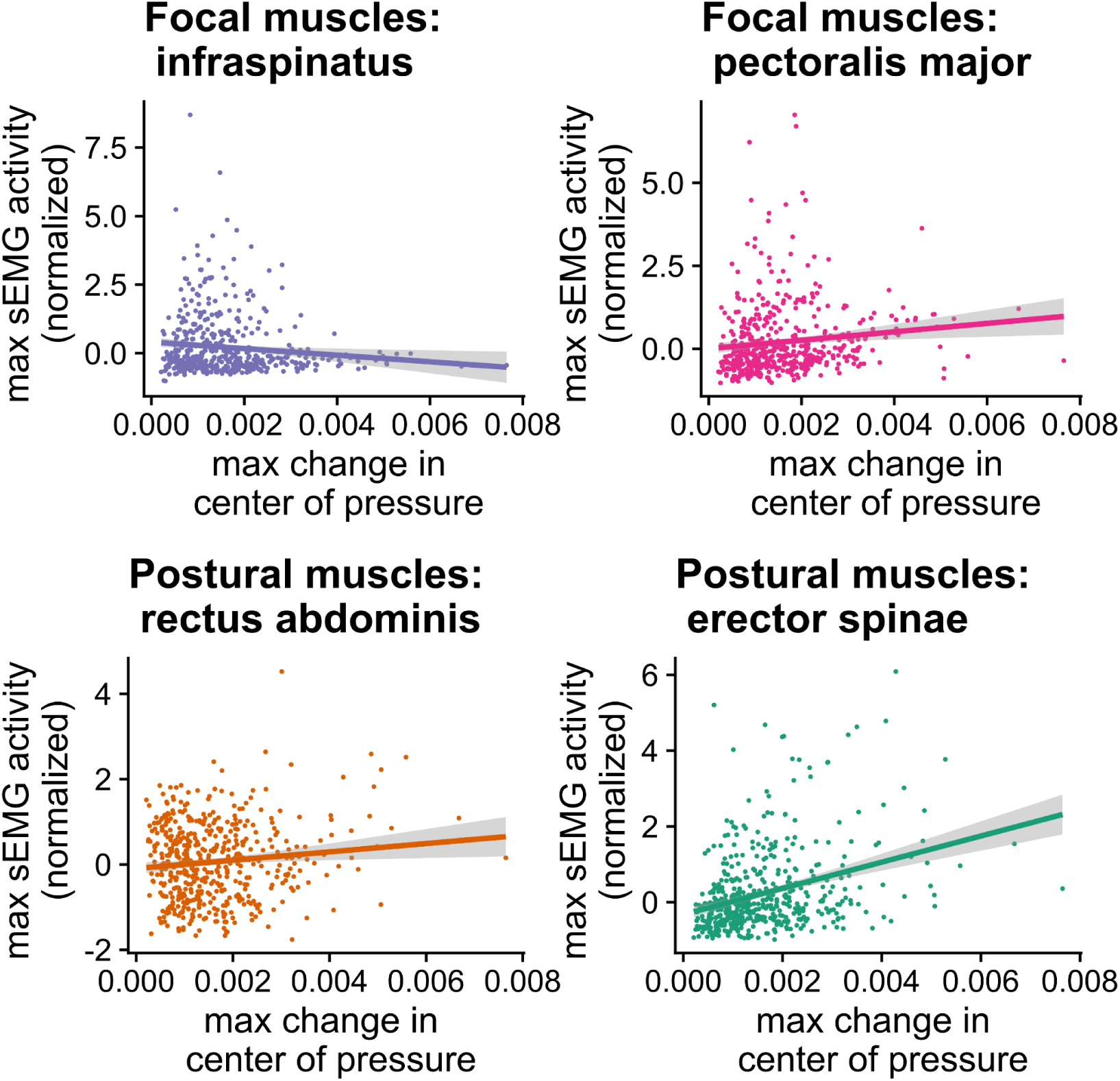
Confirmation of postural muscle activation during changes in center of mass.

**Table S12.**
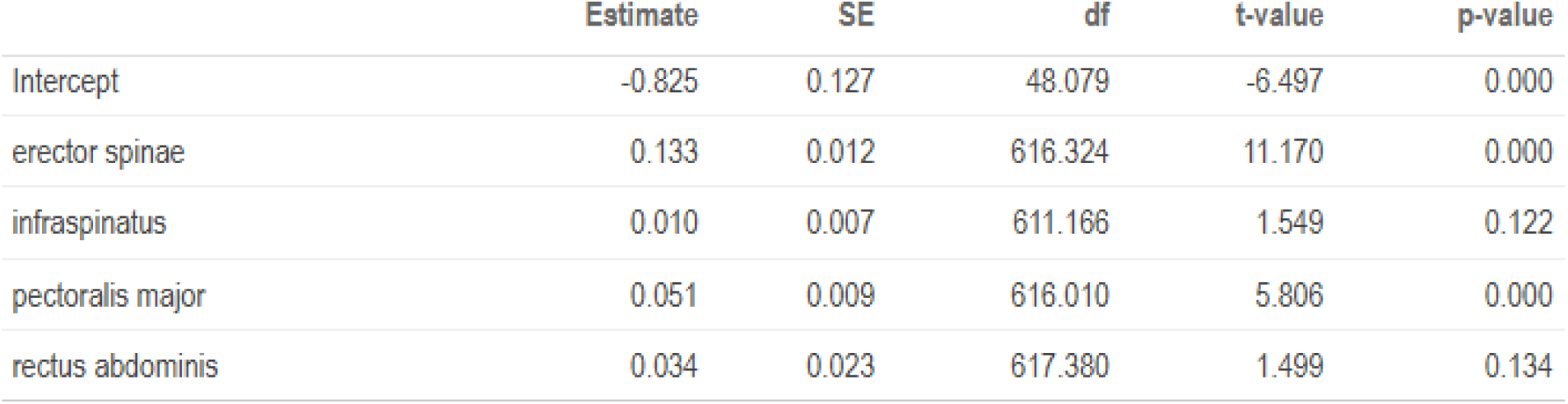
Linear mixed regression model for predicting peak change in center of pressure based on muscle activity.

|  | Estimate | SE | df | t-value | p-value |
| --- | --- | --- | --- | --- | --- |
| Intercept | -0.825 | 0.127 | 48.079 | -6.497 | 0.000 |
| erector spinae | 0.133 | 0.012 | 616.324 | 11.170 | 0.000 |
| infraspinatus | 0.010 | 0.007 | 611.166 | 1.549 | 0.122 |
| pectoralis major | 0.051 | 0.009 | 616.010 | 5.806 | 0.000 |
| rectus abdominis | 0.034 | 0.023 | 617.380 | 1.499 | 0.134 |

### Other Exploratory Analyses

Along with the confirmatory analyses reported above that relate to questions posed in the pre-registration, we report on additional, exploratory analyses. Firstly we assess for possible context-dependent interrelationships of posture, muscle activity, and voice, by producing correlation matrices per movement condition (see Tables S13, S14, S15, S16) which are represented in our synergies plot in Fig. 3 (panel F) in the main article. We see that producing different movements can recruit variable interrelationships between muscle, posture, and voice. Striking to pick out is for example that we find a direct relationship between the degree of center of pressure change and the magnitude of positive amplitude peaks in the voice.

Table S13 shows the correlation matrix for the maximum peaks in EMG (row and columns 1-4), change in center of pressure (5), and maximum vocal amplitude (6). For this table, the data only include the internal rotation condition.

**Table S13.** Correlation matrix for internal rotation.

|  | 1 | 2 | 3 | 4 | 5 |
| --- | --- | --- | --- | --- | --- |
| 1. max. EMG pectoralis major | • |  |  |  |  |
| 2. max. EMG infraspinatus | .34*** | • |  |  |  |
| 3. max. EMG rectus abdominis | -.01 | -.16 | • |  |  |
| 4. max. EMG erector spinae | .17 | .09 | .16 | • |  |
| 5. max. change in center of pressure | .28** | .17 | -.06 | .15 | • |
| 6. max. positive amplitude vocalization | .42*** | .22* | -.02 | .10 | .43*** |

Table S14 shows the correlation matrix for the maximum peaks in EMG (row and columns 1-4), change in center of pressure (5), and maximum vocal amplitude (6) For this table, the data only include the external rotation condition.

**Table S14.** Correlation matrix for external rotation.

|  | 1 | 2 | 3 | 4 | 5 |
| --- | --- | --- | --- | --- | --- |
| 1. max. EMG pectoralis major | • |  |  |  |  |
| 2. max. EMG infraspinatus | .04 | • |  |  |  |
| 3. max. EMG rectus abdominis | -.14 | .06 | • |  |  |
| 4. max. EMG erector spinae | .20* | .07 | .15 | • |  |
| 5. max. change in center of pressure | .10 | .19* | -.09 | .24** | • |
| 6. max. positive amplitude vocalization | .27** | .18* | -.05 | .25** | .11 |

Table S15 shows the correlation matrix for the maximum peaks in EMG (row and columns 1-4), change in center of pressure (5), and maximum vocal amplitude (6) For this table, the data only include the extension condition.

**Table S15:** Correlation matrix for extension.

|  | 1 | 2 | 3 | 4 | 5 |
| --- | --- | --- | --- | --- | --- |
| 1. max. EMG pectoralis major | • |  |  |  |  |
| 2. max. EMG infraspinatus | .18* | • |  |  |  |
| 3. max. EMG rectus abdominis | .06 | -.02 | • |  |  |
| 4. max. EMG erector spinae | .20* | .05 | .15 | • |  |
| 5. max. change in center of pressure | .19* | .23** | .26** | .23** | • |
| 6. max. positive amplitude vocalization | -.02 | .29*** | -.01 | .26** | .15 |

Table S16 shows the correlation matrix for the maximum peaks in EMG (row and columns 1-4), change in center of pressure (5), and maximum vocal amplitude (6) For this table, the data only include the flexion condition.

**Table S16:** Correlation matrix for extension.

|  | 1 | 2 | 3 | 4 | 5 |
| --- | --- | --- | --- | --- | --- |
| 1. max. EMG pectoralis major | • |  |  |  |  |
| 2. max. EMG infraspinatus | .18* | • |  |  |  |
| 3. max. EMG rectus abdominis | -.19* | -.02 | • |  |  |
| 4. max. EMG erector spinae | .03 | .26** | .05 | • |  |
| 5. max. change in center of pressure | -.02 | .29** | .07 | .06 | • |
| 6. max. positive amplitude vocalization | .09 | .23** | .00 | .07 | .14 |

Figure S8 visualizes the positive correlation between posture (max. change in center of pressure) and vocalization amplitude (positive peaks in amplitude) found for the internal rotation condition in Table S13.

**Figure S8.**
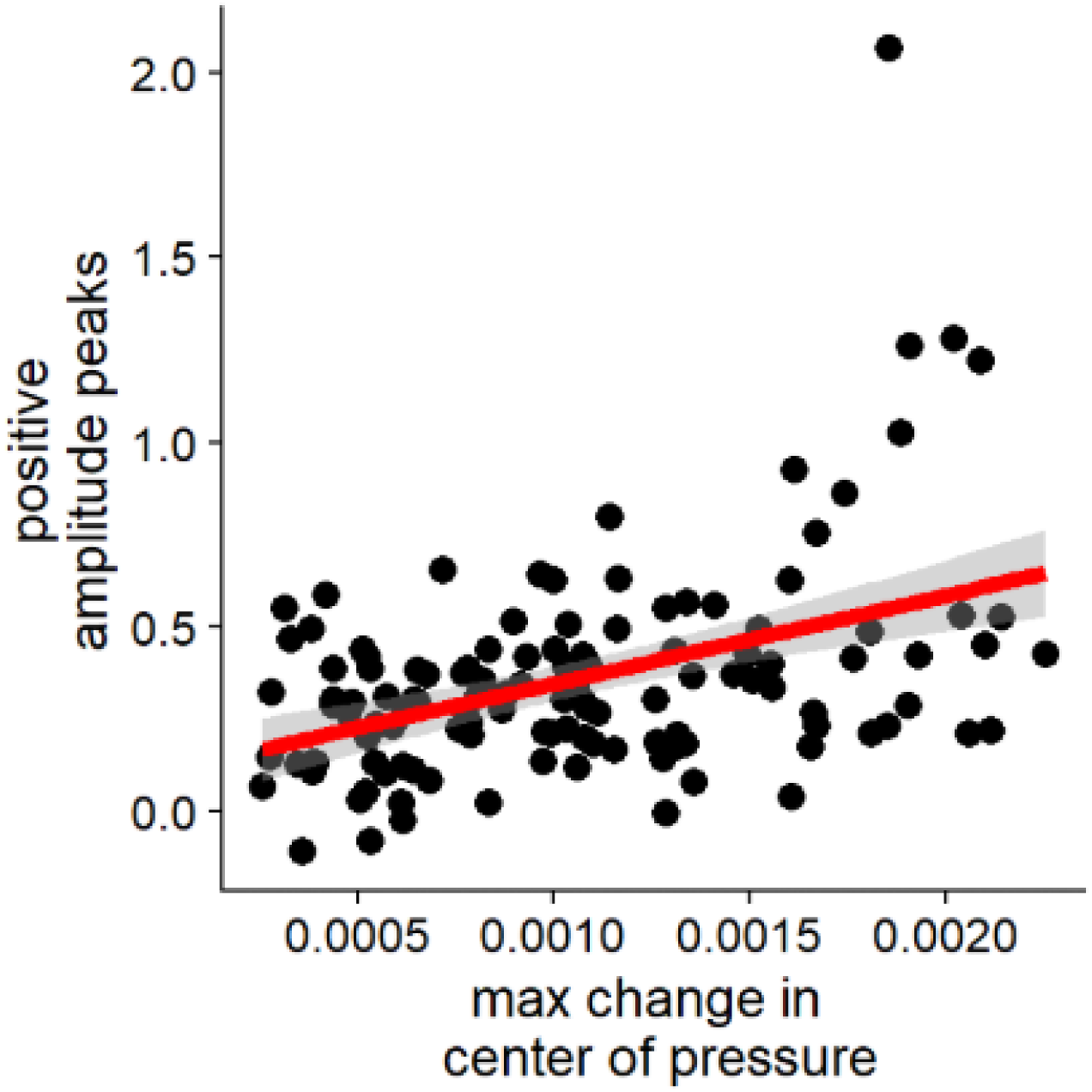
Correlation of posture and amplitude for internal rotation.

We finally explored the timing relationship of negative and positive peaks in the amplitude of the voice relative to the peak in speed of the hand. The density plot in Fig. S10 visualizes the temporal relationship between peak speed in movement and negative (left) and positive (right) peaks in the amplitude envelope, relative to the moment the moving hand reaches a peak in speed during the trial (i.e., t=0, at peak speed of the wrist). Our findings corroborate our general observation that positive (rather than negative) peaks seem to be more robustly driven by gesture kinetics. In the timing relations obtained, the negative peaks that we observe are more variably distributed around peak speed, while for the positive peaks, we see clear occurrences just around the movement onset (just before the peak speed at t = 0). This result, next to our more robust relationship between positive peaks and the muscle and posture signals, makes us conclude that in general the movements performed in our experiment are increasing subglottal pressures increasing the voice’s amplitude.

**Figure S9:**
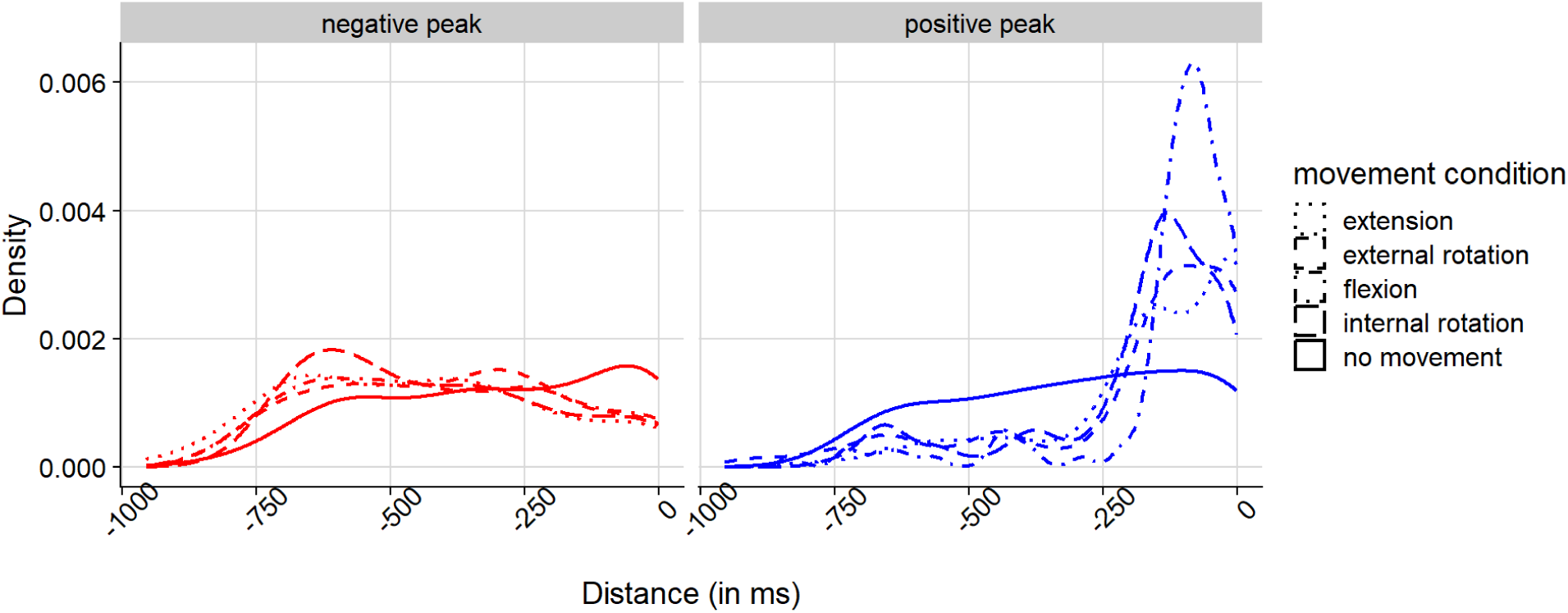
Density plot showing temporal distance (in milliseconds) of positive and negative peaks in amplitude from peak speed in wrist movement.

### Exploring potential confounds: Causality direction

Fig. S10 shows the amplitude envelope and EMG activity of an interval around the movement onset. Here we have modeled the continuous trajectories of the amplitude envelope and all the other signals (EMG’s, posture) using Generalized Additive Modeling for each movement condition, setting participants as random intercept. To reduce over-estimation of any effects, we also accounted in the model for the autocorrelation of the residuals (24, 39). Based on the predicted non-linear slopes that we generated for an interval of 600 ms we determine firstly the approximated onsets of the continuous measures. Onsets are determined as the peak in the 2nd derivative with respect to time of the GAM trajectory for a particular signal and condition. In general, we only consider the presence of onsets when the overall peak of the signal exceeds 0.75 x standard deviation from the mean (otherwise the timings reported are difficult to compare). For the figure, we thus center our data based on the predicted onset that leads to the positive peak in the amplitude envelope. In this way, we can isolate when the amplitude envelope starts to respond relative to the other signals. Firstly, we determined from the GAM trajectories the estimated onset of the vocalization peak (i.e., the moment at which the amplitude starts to rise to the peak in the voice amplitude envelope envelope), as well as the onset of an approximated negative peak, if present (i.e., the moment at which the amplitude starts to fall to the negative peak envelope). We learn from this that the minor negative peaks tend to precede the positive peak, suggesting a possible ‘anticipatory vocal adjustment’. Importantly, however, in these supplemental materials, we further show that the observed negative peaks do not happen as consistently as compared to the positive peaks (see Figure S9). This further corroborates the idea that positive peaks are directly related to physical impulses, while negative peaks are a type of correlated anticipatory/reactionary activity of the vocal system.

**Figure S10.**
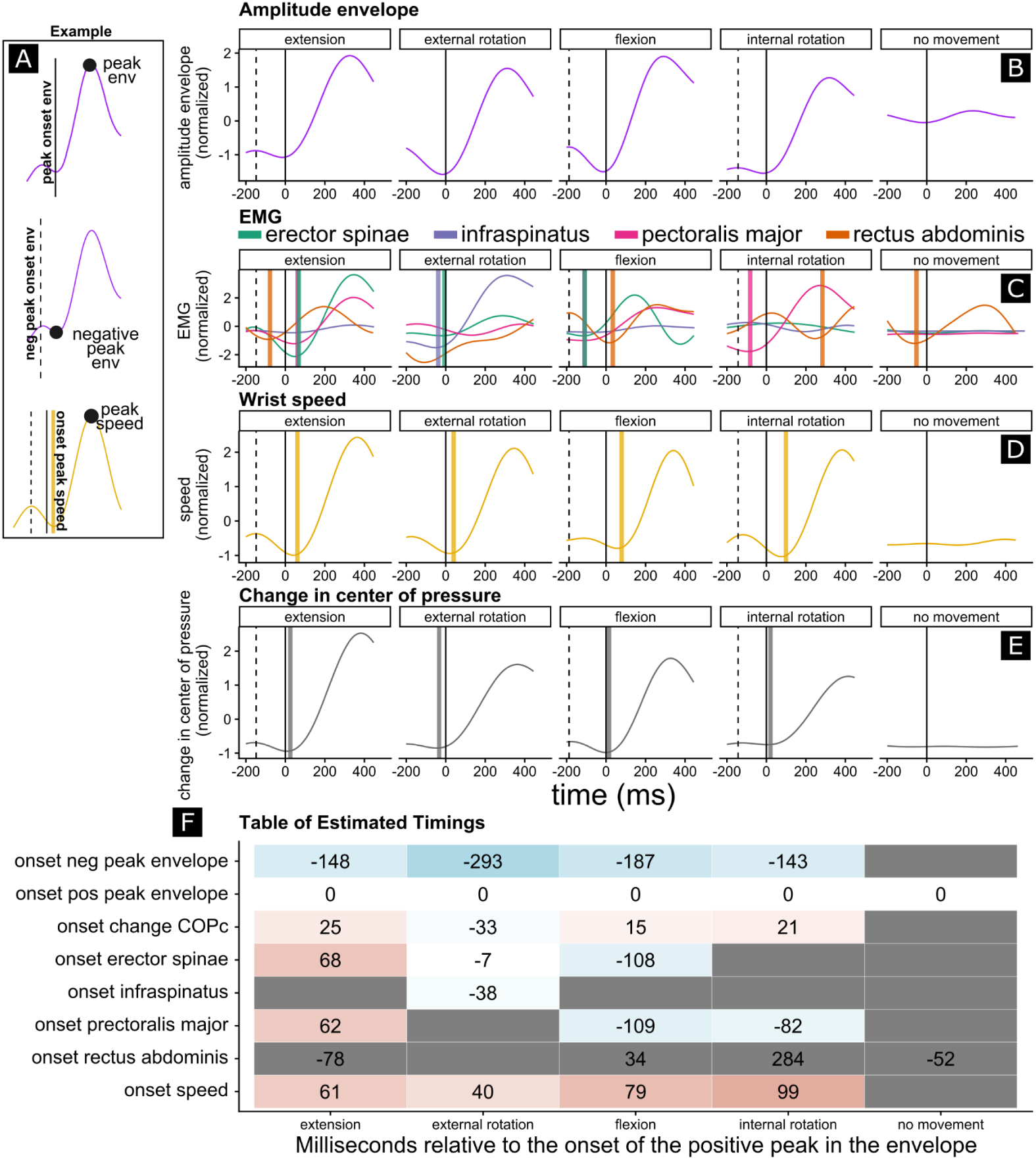
Smoothed conditional means for vocalization and muscle activity (with gam). GAMM-approximated trajectories over time, where time is centered at 0 at the vocalization onset. It can be seen that especially for the flexion movement there are clear anticipatory postural muscle activations of the erector spinae (Aruin and Latash 1995) followed by the rectus abdominis. For the extension condition, this activation pattern is reversed as one would expect given that the impulse vector should be directed in the opposite direction. Thus even though the erector abdominis proves not a very reliable site for measuring muscle activity, it still patterns in sensible ways. Note in general, that changes in vocalization are estimated to happen before observable changes in movement. Changes in vocalization are sometimes preceded by muscle activity and center of pressure changes. This yields further support that it is forces (kinetics), rather than movement (kinematics), that affects the voice.

### Exploring potential confounds: Subtle differential effects on vocalization

From our confirmatory peak analyses, we concluded that there were no reliable differences between the different movement conditions in terms of affecting the magnitude of the positive peak in vocalization. Now that we have modeled the trajectories with GAMs, we can perform a statistical test for whether the different movement-only conditions affect the vocalization amplitude envelopes in terms of their trajectory, rather than simply differing in the magnitude of the peak of the trajectory. We determine the trajectory along an interval of -500ms to +500ms relative to the onset of the arm movement (time = 0 when the movement initiates). The GAM analysis here is much more powerful as we take in millions of time series observations (as compared to the by-trial peak generation).

The GAM model with an autoregressive component, with participants as random intercepts and movement-only conditions, was statistically reliable, *F*(1818052.92, 1818025.83) = 170.97, *p* < .0001. The smooth term statistics in Table S17 show that there was a general nonlinearity to the amplitude envelope that characterized all movement-only conditions, as evidenced by the statistically reliable smooth for time. Additionally, the flexion, external rotation, and internal rotation, each had a movement-specific pattern that was statistically reliable. This means that there are statistically robust deviations in the shaping of the amplitude envelope trajectories for different types of movement (except for the extension condition).

**Table S17.** Tests of smooth terms in the model.

|  | Term | edf | Ref_df | F_stat | p_value |
| --- | --- | --- | --- | --- | --- |
| s(movstart) | s(movstart) | 7.99 | 8.00 | 8.18 | < 2.2e-16 |
| s(movstart):movement_conditionextension_stop | s(movstart):movement_conditionextension_stop | 6.22 | 6.25 | 2.43 | 0.0157260 |
| s(movstart):movement_conditionexternal_rotation_stop | s(movstart):movement_conditionexternal_rotation_stop | 7.22 | 7.25 | 3.04 | 0.0075828 |
| s(movstart):movement_conditionflexion_stop | s(movstart):movement_conditionflexion_stop | 7.20 | 7.25 | 2.47 | 0.0080897 |
| s(movstart):movement_conditioninternal_rotation_stop | s(movstart):movement_conditioninternal_rotation_stop | 7.22 | 7.25 | 4.31 | 3.525e-05 |
| s(pp) | s(pp) | 0.92 | 1.00 | 11.86 | 0.0003326 |

**Figure S11.**
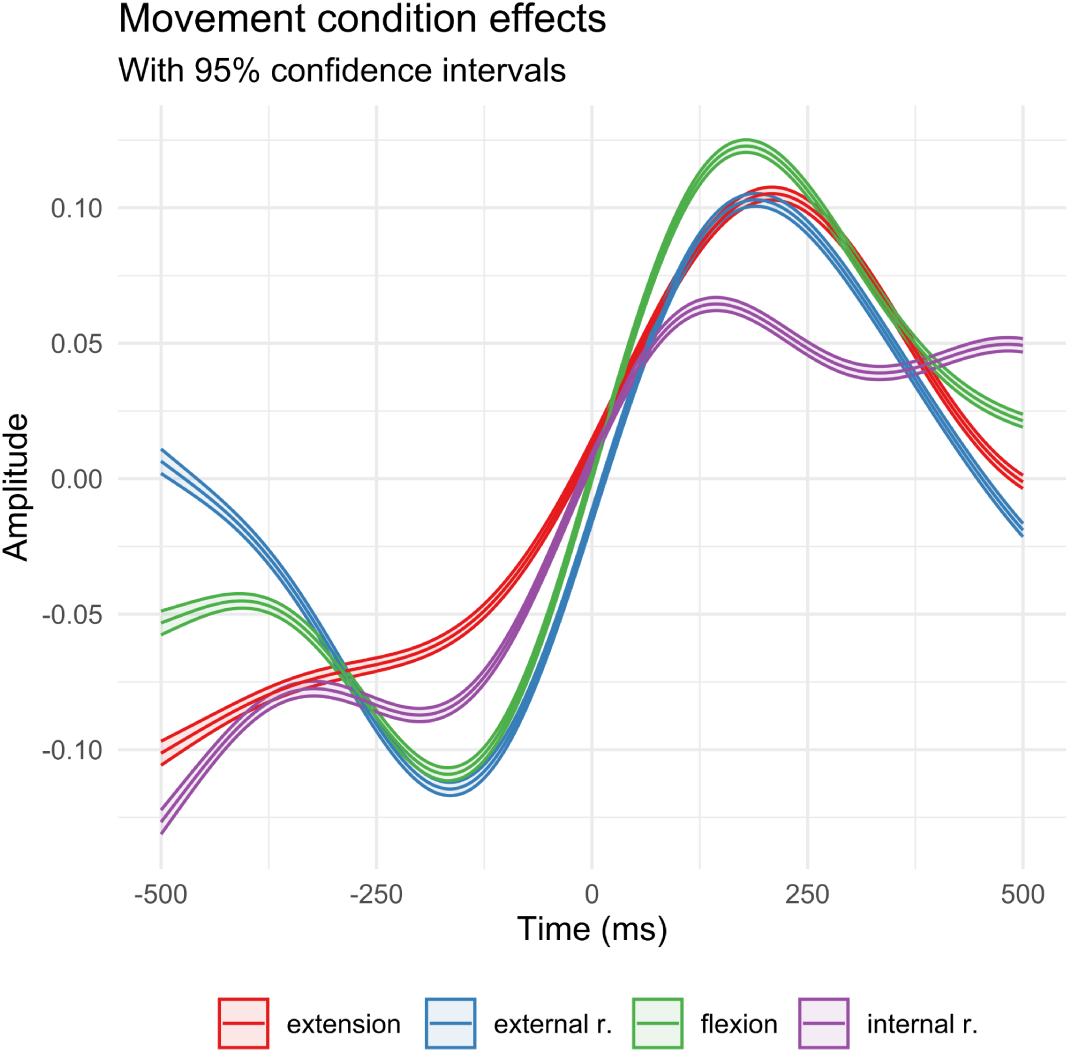
GAM-approximated vocal trajectories for movement-only conditions with confidence intervals.

**Table S18.**
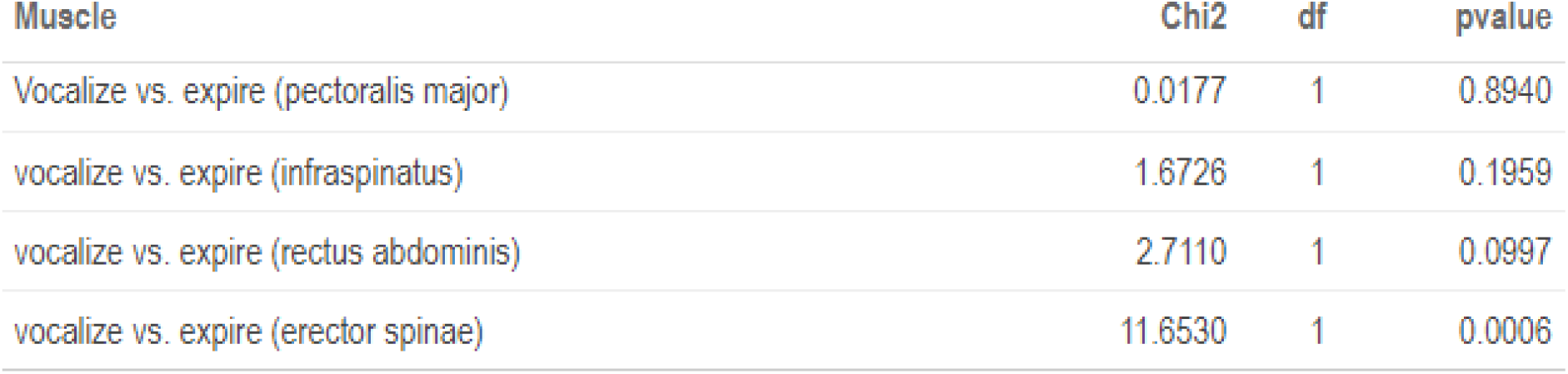
Muscle activity predicted by presence vocalization versus expiration for the movement only conditions (testing against a model accounting for weight and movement conditions).

| Muscle | Chi2 | df | pvalue |
| --- | --- | --- | --- |
| Vocalize vs. expire (pectoralis major) | 0.0177 | 1 | 0.8940 |
| vocalize vs. expire (infraspinatus) | 1.6726 | 1 | 0.1959 |
| vocalize vs. expire (rectus abdominis) | 2.7110 | 1 | 0.0997 |
| vocalize vs. expire (erector spinae) | 11.6530 | 1 | 0.0006 |

### Exploring potential confounds: Muscle activity for the different movement and vocalization conditions

Importantly, it is possible that vocalizing recruits more muscle activity in general, including the no movement condition, because the subglottal pressures are higher for vocalization due to increased air resistance of the adducted vocal folds as compared to the partially closed lips for controlled expiration. To test an overall increased erector spinae activity, rather than a differentiated increase for movement trials only, we first assessed whether a model predicting erector spinae activity for all movement conditions (including the no movement condition), over and above movement presence and weight conditions, was a better model than a comparison model with only movement and weight condition. This model increased explained variance, χ2(1) = 9.95, *p* = 0.0016, and indeed in general there was an overall increased effect of vocalization on erector spinae activity, b = 0.452, t(1232.82) = 3.155, *p* = 0.0016. Since another model with an added interaction term of movement presence and vocal condition did not reliably improve relative to a model with simple effects, χ2(1) = 1.427, *p* = 0.2322, we conclude that in general vocalizations recruit higher erector spinae activity than expiration in these tasks. This thereby refutes the possibility that there is increased muscle activation in anticipation of movement perturbing the voice and thereby appearing as if arm-movement directly affects the voice. After all, in the no movement condition there is also a higher erector spinae recruitment when vocalizing vs. expiring.

**Figure S11.**
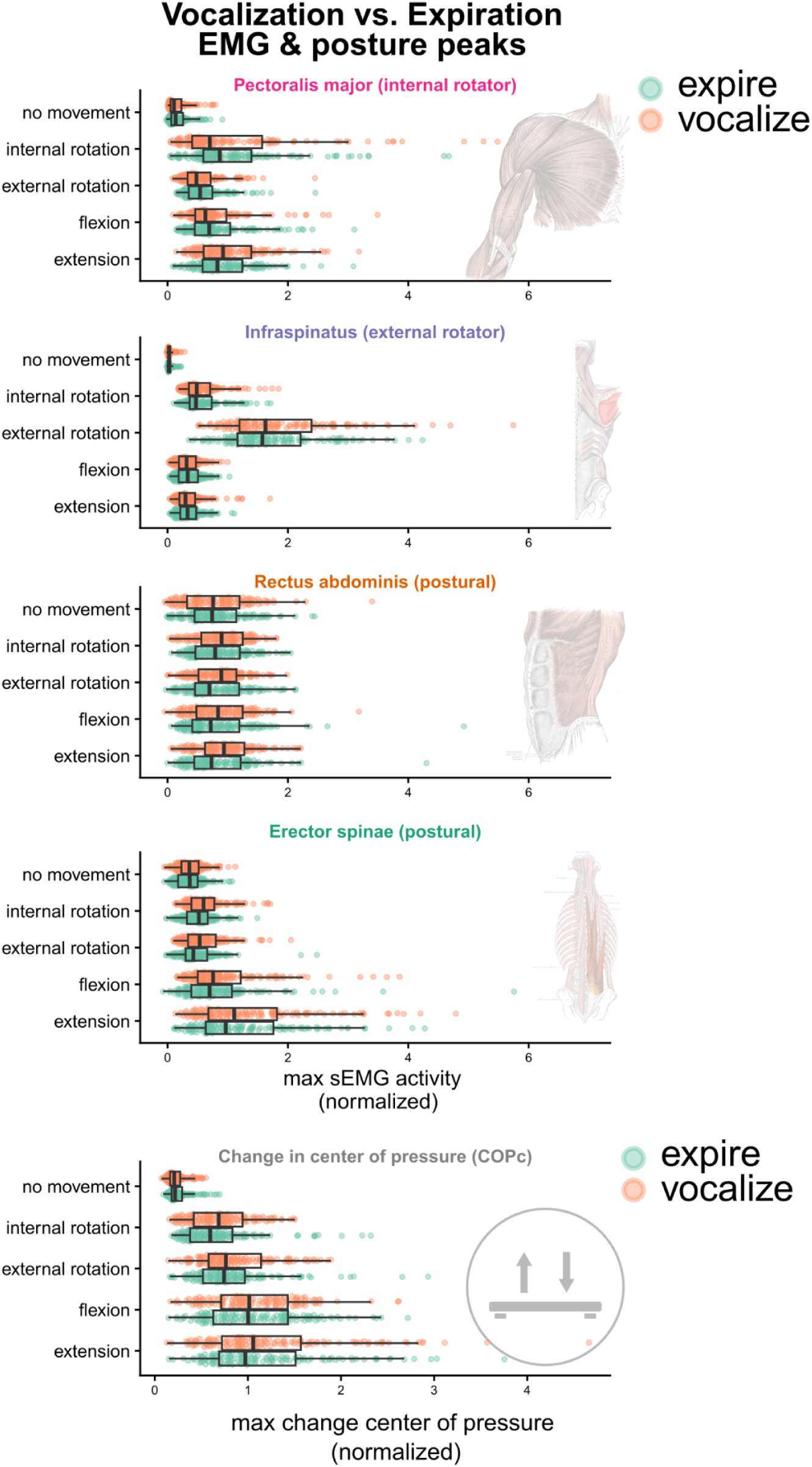
Muscle and postural activations for vocalization versus expiration.

**Table S19.** Normalized peak muscle activity for the different movement and vocalization conditions. These are the numerical results associated with Figure S4. The values indicate the peak EMG activity normalized for each muscle.

| Muscle | Movement condition | Vocal condition | Mean | SD | Low | High |
| --- | --- | --- | --- | --- | --- | --- |
|  |  |  |  |  | 95% CI | 95% CI |
| pectoralis major | no movement | expire | 0.18 | 0.17 | 0.15 | 0.21 |
| pectoralis major | no movement | vocalize | 0.17 | 0.17 | 0.14 | 0.19 |
| pectoralis major | internal rotation | expire | 1.14 | 0.87 | 0.99 | 1.29 |
| pectoralis major | internal rotation | vocalize | 1.12 | 1.09 | 0.93 | 1.31 |
| pectoralis major | external rotation | expire | 0.60 | 0.36 | 0.54 | 0.67 |
| pectoralis major | external rotation | vocalize | 0.58 | 0.36 | 0.51 | 0.64 |
| pectoralis major | flexion | expire | 0.83 | 0.54 | 0.73 | 0.92 |
| pectoralis major | flexion | vocalize | 0.79 | 0.54 | 0.69 | 0.89 |
| pectoralis major | extension | expire | 0.95 | 0.52 | 0.86 | 1.04 |
| pectoralis major | extension | vocalize | 1.04 | 0.57 | 0.94 | 1.14 |
| infraspinus | no movement | expire | 0.04 | 0.04 | 0.03 | 0.05 |
| infraspinus | no movement | vocalize | 0.04 | 0.05 | 0.03 | 0.05 |
| infraspinus | internal rotation | expire | 0.59 | 0.33 | 0.53 | 0.65 |
| infraspinus | internal rotation | vocalize | 0.58 | 0.33 | 0.52 | 0.63 |
| infraspinus | external rotation | expire | 1.70 | 0.85 | 1.55 | 1.85 |
| infraspinus | external rotation | vocalize | 1.88 | 1.07 | 1.69 | 2.07 |
| infraspinus | flexion | expire | 0.35 | 0.21 | 0.32 | 0.39 |
| infraspinus | flexion | vocalize | 0.34 | 0.20 | 0.30 | 0.37 |
| infraspinus | extension | expire | 0.37 | 0.21 | 0.33 | 0.41 |
| infraspinus | extension | vocalize | 0.37 | 0.26 | 0.33 | 0.42 |
| rectus abdominis | no movement | expire | 0.81 | 0.51 | 0.72 | 0.90 |
| rectus abdominis | no movement | vocalize | 0.81 | 0.57 | 0.71 | 0.91 |
| rectus abdominis | internal rotation | expire | 0.85 | 0.45 | 0.77 | 0.93 |
| rectus abdominis | internal rotation | vocalize | 0.89 | 0.43 | 0.81 | 0.96 |
| rectus abdominis | external rotation | expire | 0.83 | 0.51 | 0.74 | 0.92 |
| rectus abdominis | external rotation | vocalize | 0.85 | 0.42 | 0.78 | 0.93 |
| rectus abdominis | flexion | expire | 0.86 | 0.63 | 0.75 | 0.97 |
| rectus abdominis | flexion | vocalize | 0.88 | 0.51 | 0.79 | 0.97 |
| rectus abdominis | extension | expire | 0.86 | 0.58 | 0.76 | 0.96 |
| rectus abdominis | extension | vocalize | 0.94 | 0.49 | 0.86 | 1.03 |
| erector spinae | no movement | expire | 0.37 | 0.23 | 0.33 | 0.41 |
| erector spinae | no movement | vocalize | 0.39 | 0.22 | 0.35 | 0.43 |
| erector spinae | internal rotation | expire | 0.52 | 0.27 | 0.47 | 0.57 |
| erector spinae | internal rotation | vocalize | 0.64 | 0.34 | 0.58 | 0.70 |
| erector spinae | external rotation | expire | 0.50 | 0.35 | 0.44 | 0.56 |
| erector spinae | external rotation | vocalize | 0.60 | 0.36 | 0.54 | 0.67 |
| erector spinae | flexion | expire | 0.85 | 0.74 | 0.72 | 0.98 |
| erector spinae | flexion | vocalize | 0.97 | 0.71 | 0.84 | 1.09 |
| erector spinae | extension | expire | 1.24 | 0.88 | 1.08 | 1.39 |
| erector spinae | extension | vocalize | 1.39 | 0.97 | 1.22 | 1.56 |

## Author note

Some of the text in this report is copied verbatim from the pre-registration as well as the supporting information. Findings of this report concerning research question 1 were reported in <u>Proceedings</u> of the International Conference on the Evolution of Language (EVOLANG2024).

## Acknowledgements

WP is funded by a VENI grant (VI.Veni 0.201G.047: PI Wim Pouw) and by the Language Interaction Consortium. We would like to thank Pascal de Water for the technical support in implementing LabStreamingLayer to ensure synchronized signal recording and his general support for building the lab that accommodated this research. We would also like to thank the participants who volunteered in this study. We further want to thank four reviewers for there very valuable and detailed comments that has improved the report considerably.

## Data Accessibility

There is an <u>Rmarkdown notebook</u> with code alongside the methods and results, providing a fully computationally reproducible pre-processing and analyses procedure. The full dataset with masked videos to reduce identity exposure will be openly available on the Donders Repository. Currently the full dataset is only downloadable for reviewers with a password ‘kineticspouw’, by following the link in the <u>dataset folder</u> where the reproducible code is linked to https://github.com/WimPouw/kineticsvoice/.

## Footnotes

1 This pre-registration received a timestamp on 30-5-2023 and that version was then freezed and became immutable.

2 Note we ignored the interaction of movement and weight, and planned to assess the effect of weight over and above the movement conditions in our pre-registration. This analysis ignores the fact that wearing a weight should have no effect when there is no movement. This planned analysis is therefore not strictly assessing vocal effects of wearing weights for upper limb movement. This planned analysis, which first accounts for variability in movement conditions but only assesses a main effect of weights, shows an overall effect of weight for positive peaks (*p* = .005), while conclusions for negative peaks remain unchanged (*p* = .024). We report a more conservative analysis here however, by simply assessing the effect of weight vs. no weight on vocal peaks in the movement-only conditions.

## References

1. J. A. Seikel, D. G. Drumright, D. J. Hudock, Anatomy & Physiology for Speech, Language, and Hearing (Plural Publishing, Incorporated, 2019).

2. P. W. Hodges, S. C. Gandevia, Changes in intra-abdominal pressure during postural and respiratory activation of the human diaphragm. J. Appl. Physiol. 89, 967–976 (2000).

3. P. W. Hodges, C. A. Richardson, Feedforward contraction of transversus abdominis is not influenced by the direction of arm movement. Exp Brain Res 114, 362–370 (1997).

4. A. S. Aruin, M. L. Latash, Directional specificity of postural muscles in feed-forward postural reactions during fast voluntary arm movements. Exp Brain Res 103, 323–332 (1995).

5. S. L. Morris, B. Lay, G. T. Allison, Transversus abdominis is part of a global not local muscle synergy during arm movement. Hum Mov Sci 32, 1176–1185 (2013).

6. D. Lasserson, et al., Differences in motor activation of voluntary and reflex cough in humans. Thorax 61, 699–705 (2006).

7. V. Pettersen, R. H. Westgaard, Muscle activity in professional classical singing: a study on muscles in the shoulder, neck and trunk. Logopedics Phoniatrics Vocology 29, 56–65 (2004).

8. V. Pettersen, K. Bjørkøy, H. Torp, R. H. Westgaard, Neck and shoulder muscle activity and thorax movement in singing and speaking tasks with variation in vocal loudness and pitch. J Voice 19, 623–634 (2005).

9. T. E. Dolmage, et al., Arm elevation and coordinated breathing strategies in patients with COPD. Chest 144, 128–135 (2013).

10. X. Liu, et al., Evaluation of isokinetic muscle strength of upper limb and the relationship with pulmonary function and respiratory muscle strength in stable COPD patients. COPD 14, 2027–2036 (2019).

11. M. Zedka, A. Prochazka, Phasic activity in the human erector spinae during repetitive hand movements. J Physiol 504, 727–734 (1997).

12. . S. Fuchs, A. Rochet-Capellan, The Respiratory Foundations of Spoken Language. Annual Review of Linguistics 7, null (2021).

13. P. Wagner, A. Ćwiek, B. Samlowski, Exploiting the speech-gesture link to capture fine-grained prosodic prominence impressions and listening strategies. Journal of Phonetics 76, 100911 (2019).

14. J. Krivokapić, Gestural coordination at prosodic boundaries and its role for prosodic structure and speech planning processes. Philos Trans R Soc Lond B Biol Sci 369, 1–44 (2014).

15. D. Bolinger, Intonation and gesture. American Speech 58, 156–174 (1983).

16. W. Pouw, S. Fuchs, Origins of vocal-entangled gesture. Neuroscience & Biobehavioral Reviews 141, 104836 (2022).

17. T. Jenkins, W. Pouw, Gesture–speech coupling in persons with aphasia: A kinematic-acoustic analysis. Journal of Experimental Psychology: General 152, 1469–1483 (2023).

18. L. Pearson, W. Pouw, Gesture–vocal coupling in Karnatak music performance: A neuro–bodily distributed aesthetic entanglement. Annals of the New York Academy of Sciences n/a (2022).

19. W. Pouw, A. Paxton, S. J. Harrison, J. A. Dixon, Acoustic information about upper limb movement in voicing. PNAS 117, 11364–11367 (2020).

20. W. Pouw, L. de Jonge-Hoekstra, S. J. Harrison, A. Paxton, J. A. Dixon, Gesture-speech physics in fluent speech and rhythmic upper limb movements. Annals of the New York Academy of Sciences 1491, 89–105 (2020).

21. W. Pouw, S. J. Harrison, N. Esteve-Gibert, J. A. Dixon, Energy flows in gesture-speech physics: The respiratory-vocal system and its coupling with hand gestures. The Journal of the Acoustical Society of America 148, 1231–1247 (2020).

22. W. Pouw, S. J. Harrison, J. A. Dixon, The importance of visual control and biomechanics in the regulation of gesture-speech synchrony for an individual deprived of proprioceptive feedback of body position. Scientific Reports (2020). 10.31234/osf.io/7g3es.

23. W. Pouw, A. Paxton, S. J. Harrison, J. A. Dixon, Acoustic specification of upper limb movement in voicing in *Proceedings of the 6th Meeting of Gesture and Speech in Interaction*, (Universitaetsbibliothek Paderborn, 2019), pp. 75–80.

24. H. Serré, M. Dohen, S. Fuchs, S. Gerber, A. Rochet-Capellan, Leg movements affect speech intensity. Journal of Neurophysiology (2022). 10.1152/jn.00282.2022.

25. S. Harrison, ‘This you may NNNNNNEVER have heard before’: initial lengthening of accented negative items as vocal-entangled gestures. Language and Cognition 16, 1778–1811 (2024).

26. W. Pouw, S. J. Harrison, J. A. Dixon, Gesture-speech physics: The biomechanical basis of the emergence of gesture-speech synchrony. Journal of Experimental Psychology: General 149, 391–404 (2019).

27. S. Nadkarni, P. Rao, M. Clayton, Identifying Melodic Motifs and Stable Notes from Gestural Information in Indian Vocal Performances. Transactions of the International Society for Music Information Retrieval 7, 246–263 (2024).

28. R. Werner, L. Selen, W. Pouw, Arm movements increase acoustic markers of expiratory flow in Proceedings of the International Conference of Speech Prosody (SpeechProsody2024), (2024).

29. P. L. Rohrer, Y. Hong, H. R. Bosker, Gestures time to vowel onset and change the acoustics of the word in Mandarin in (2024), pp. 866–870.

30. P. W. Hodges, R. Sapsford, L. H. M. Pengel, Postural and respiratory functions of the pelvic floor muscles. Neurourol. Urodyn. 26, 362–371 (2007).

31. V. Pettersen, Preliminary findings on the classical singer’s use of the pectoralis major muscle. FPL 58, 427–439 (2006).

32. P. Cavallari, F. Bolzoni, C. Bruttini, R. Esposti, The Organization and Control of Intra-Limb Anticipatory Postural Adjustments and Their Role in Movement Performance. Front. Hum. Neurosci. 10 (2016).

33. S. Bouisset, M.-C. Do, Posture, dynamic stability, and voluntary movement. Neurophysiologie Clinique/Clinical Neurophysiology 38, 345–362 (2008).

34. F. G. Baldissera, L. Tesio, APAs Constraints to Voluntary Movements: The Case for Limb Movements Coupling. Frontiers in Human Neuroscience 11 (2017).

35. W. C. Lancaster, O. W. Henson, A. W. Keating, Respiratory muscle activity in relation to vocalization in flying bats. Journal of Experimental Biology 198, 175–191 (1995).

36. B. G. Cooper, F. Goller, Multimodal signals: enhancement and constraint of song motor patterns by visual display. Science 303, 544–546 (2004).

37. M. T. Turvey, S. T. Fonseca, The medium of haptic perception: A tensegrity hypothesis. Journal of Motor Behavior 46, 143–187 (2014).

38. M. Brysbaert, M. Stevens, Power Analysis and Effect Size in Mixed Effects Models: A Tutorial. Journal of Cognition 1, 9 (2018).

39. M. Wieling, Analyzing dynamic phonetic data using generalized additive mixed modeling: A tutorial focusing on articulatory differences between L1 and L2 speakers of English. Journal of Phonetics 70, 86–116 (2018).

40. T. J. Roberts, A. M. Gabaldón, Interpreting muscle function from EMG: lessons learned from direct measurements of muscle force. Integrative and Comparative Biology 48, 312–320 (2008).

41. B. Parrell, L. Goldstein, S. Lee, D. Byrd, Spatiotemporal coupling between speech and manual motor actions. Journal of Phonetics 42, 1–11 (2014).

42. G. Zelic, J. Kim, C. Davis, Articulatory constraints on spontaneous entrainment between speech and manual gesture. Human Movement Science 42, 232–245 (2015).

43. W. Pouw, J. A. Dixon, Entrainment and modulation of gesture–speech synchrony under delayed auditory feedback. Cognitive Science 43, e12721 (2019).

44. P. J. Treffner, M. Peter, Intentional and attentional dynamics of speech–hand coordination. Human Movement Science 21, 641–697 (2002).

45. K. Garvin, E. Spradling, K. Franich, Co-speech gestures influence the magnitude and stability of articulatory movements: Evidence for coupling-based enhancement. [Preprint] (2024). Available at: https://www.researchsquare.com/article/rs-5073434/v1 [Accessed 15 December 2024].

46. S. P. Coundouris, A. G. Adams, S. A. Grainger, J. D. Henry, Social perceptual function in Parkinson’s disease: A meta-analysis. Neuroscience and Biobehavioral Reviews 104, 255–267 (2019).

47. A. K. Ho, R. Iansek, J. L. Bradshaw, Motor Instability in Parkinsonian Speech Intensity. Cognitive and Behavioral Neurology 14, 109–116 (2001).

48. S. Sharma, K. Fleck, S. Winslow, K. Rothermich, The Impact of Parkinson’s Disease on Social Communication: An Exploratory Questionnaire Study. [Preprint] (2021). Available at: https://osf.io/preprints/socarxiv/3x42a/ [Accessed 24 August 2022].

49. M. J. Richardson, K. L. Marsh, R. C. Schmidt, Effects of visual and verbal interaction on unintentional interpersonal coordination. J Exp Psychol Hum Percept Perform 31, 62–79 (2005).

50. J. J. Gibson, The senses considered as perceptual systems (Houghton Mifflin, 1966).

51. P. J. Thibault, Distributed languaging, affective dynamics, and the human ecology: The sense-making body, Vol. 1 (Routledge/Taylor & Francis Group, 2021).

52. N. A. Miller, et al., The effects of humming and pitch on craniofacial and craniocervical morphology measured using MRI. Journal of Voice 26, 90–101 (2012).

53. N. A. Miller, J. S. Gregory, R. M. Aspden, P. J. Stollery, F. J. Gilbert, Using active shape modeling based on MRI to study morphologic and pitch-related functional changes affecting vocal structures and the airway. Journal of Voice 28, 554–564 (2014).

54. S. Leonetti, A. Ravignani, W. Pouw, A cross-species framework for classifying sound-movement couplings. Neurosci Biobehav Rev 167, 105911 (2024).

55. D. F. Boggs, Interactions between locomotion and ventilation in tetrapods. Comparative Biochemistry and Physiology Part A: Molecular & Integrative Physiology 133, 269–288 (2002).

56. C. Lugaresi, et al., MediaPipe: A Framework for Building Perception Pipelines. [Preprint] (2019). Available at: http://arxiv.org/abs/1906.08172 [Accessed 30 August 2022].

57. B. Owoyele, J. Trujillo, G. de Melo, W. Pouw, Masked-piper: Masking personal identities in visual recordings while preserving multimodal information. SoftwareX [Preprint] (2022). Available at: https://psyarxiv.com/bpt26/ [Accessed 3 June 2022].

58. R. V. Lenth, et al., emmeans: Estimated Marginal Means, aka Least-Squares Means. (2021). Deposited 21 March 2021.

